# *On-chip* perivascular niche with patient-derived glioma cells

**DOI:** 10.1101/2020.12.23.424179

**Authors:** Magda Gerigk, Harry Bulstrode, HaoTian Harvey Shi, Felix Tönisen, Camilla Cerutti, Gillian Morrison, David Rowitch, Yan Yan Shery Huang

## Abstract

Glioblastoma multiforme (GBM), is the most common and the most aggressive type of primary brain malignancy. Glioblastoma stem-*like* cells (GSCs) are able to migrate in vascular niches within or away from the tumour mass, increasing tumour resistance to patient treatments and contributing to relapses. To study individual GSCs migration and their interactions with the microenvironment in the vasculature, there is a need to develop a model of human blood vessels *in vitro*. Herein, we report a systematic study on the interaction between patient-derived glioma stem-*like* cell lines with different organotypic perivascular niche models. A microfluidic chip integrated with an extracellular matrix was fabricated to support the culture of rounded microvessels, formed with endothelial cells from three different organs, (1) human brain microvascular endothelial cells (hCMEC/D3), (2) human umbilical vein endothelial cells (HUVECs) and, (3) human lung microvascular endothelial cells (HMVEC-L). Three-dimensional (3D) cell culture retains selected adherent and tight junction markers of the endothelial cells, and the stemness-related genes of GSCs. We optimized the experimental protocol to perform qPCR, and western blot on the co-cultured GSCs with endothelial cells forming microvessels. Endpoint biological assays showed upregulation of neovascularization-related genes in endothelial cells (e.g., angiopoietins, vascular endothelial growth factor receptors) resulted after their co-culture with GBM cells. Moreover, we measured cancer cell speed and polarization during migration towards the endothelial cell formed vessel by live-cell imaging showing that organotypic (brain cancer cells – brain endothelial microvessel) interactions differ from those within non-tissue specific vascular niches. The development and optimization of this 3D microfluidic device could provide the next level of complexity of an *in vitro* system to study the influence of glioma cells on normal brain endothelium. More importantly, it enables the possibility to conduct comparative studies to dissect the influence of 3D culture, microvessel architecture and organotypic vessel types on glioma cells’ stemness and migration.

## 1 Introduction

Glioblastoma multiforme (GBM) is the most common and aggressive form of primary brain tumour associated with poor survival. Following diagnosis of a GBM, current standard of care – surgical resection, chemotherapy, and radiation – together yield a median patient survival of less than 15 months ^1^. Although standard treatments can modestly extend survival, they fail to address molecular inter- and intra-tumour heterogeneity of GBM cells, as well as the dynamic regulation of the brain microenvironment (TME) that actively supports tumour progression and evolving treatment resistance. Moreover, tumours exhibit inherent chemo-resistance, which has been attributed to a subpopulation of cancer cells termed GBM stem-*like* cells (GSCs) ^2^. Additionally, glioblastomas can often grow and progress without angiogenesis (formation of new blood vessels) and thus escape anti-angiogenic therapies^3^. Neovascularization has long been implicated as a key feature of glioblastoma progression, and could be achieved through different mechanisms, including vascular co-option, angiogenesis, vasculogenesis, vascular mimicry, and glioblastoma-endothelial cell transdifferentiation^4–6^. Therefore, the fabrication of a 3D *in vitro* model to study the vasculature in the presence of brain cancer cells could potentially aid the discovery of new strategies to target GBM neovascularization.

Substantial progress has been made recently to develop *in vitro* 3D GBM culture models through organ-on-a-chip platforms^7,8^. For example, several microfluidic models have been developed to study brain cancer cells interfacing one or more elements of the TME (Fig S1). Ayuso *et al*. have created a versatile microfluidics platform that allowed real-time monitoring of oxygen and glucose levels within the device to study the effects on brain cancer cell proliferation^9^. In another study, Ayuso *et al*. focused on mimicking the pseudopalisade formation in the GBM microenvironment *via* nutrient starvation ^10^. They have shown that the formation of pseudopalisades stimulated more aggressive behaviours in the GBM cells. Ma *et al*. recreated a coherent GBM microenvironment by means of embedding multicellular spheroids within collagen matrix with the recreation of flow characteristics through culture perfusion ^11^. This device allowed the study of the effect of various anti-invasion drugs on the growth of GBM. In all aforementioned cases, only lateral microchannels were used without actual formation of the microvessels. Recently, examples of microfluidic models that involve microvessel representations were presented by Truong, who recreated a vascular system utilizing human umbilical vein endothelial cells (HUVECs), to study the interactions between the vasculature and the patient specific GSCs ^12^. Furthermore, building on the established process of microvasculature formation using microfluidic systems, Xiao *et al*. presented an *in vitro* model that allows the evaluation of individual GBM cells directly extracted from patients correlating to specific treatment options^13^. However, in previous studies, only HUVECs were used to form the microvessels, which are not derived from the brain, and therefore may not represent an organotypic GBM-microvasculature crosstalk. Considering the tumour microenvironment in each type of cancer, the regulation of vascular niches is likely to be organ-specific. To evaluate the impact of endothelial cells of different organ origins on GSCs phenotypes, we perform systematic studies on the interaction between patient-derived glioma stem-*like* (GSC) cell lines and perfusable microvessels generated from human microvascular endothelial cells (hCMEC/D3), human umbilical vein endothelial cells (HUVECs) and human lung microvascular endothelial cells (HMVEC-L). Although more complex 3D brain vasculature mimicking blood-brain barrier characteristics or 3D organoid models have been demonstrated, the focus of our work is to investigate fundamentally, to what extent (1) assay format, and (2) tissue specific vessel-glioma interaction, matter for creating a perivascular niche for studying GBM ^14–18^. Validation to the above questions could hold practical importance towards designing on-chip devices for glioma drug testing applications, balancing assay fidelity *versus* process simplicity and costs.

With the above goal, a simple microvessel-on-chip platform reported in our previous work ^19^ was adapted to support glioma-vessel formation. We describe the model design, the biological validation, and the use of the model as a microfluidic platform as a tool to study the cellular behaviour and properties of vascular niches and within them. Cell-cell and cell-microenvironment interactions was measured by changes in gene expression in the endothelial cells forming the microvessels. Live-cell imaging enables the tracking of dynamic events in cell culture, quantifying cancer cell velocity and polarization in the presence of a microvessel in the 3D co-culture setup. The protocols developed created the opportunity to compare the effects of interfaces between different GSCs and microvessels of various tissue origin, showing that organotypic (brain cancer cells – brain endothelial microvessel) interactions indeed differ from those tracked within non-tissue specific vascular niches (i.e. brain cancer cells – endothelium from lung or umbilical cord). This work may pave the way for developing new *in vitro* models for testing candidate drugs for future glioblastoma treatment.

## 2 Materials and Methods

### 2.1 Microfluidics device fabrication

The microfluidic device used for the construction of the microvessel-on-a-chip experiments was detailed in Fig 1 ^19^. Briefly, the microfluidic device was designed using AutoCAD software. The design contains two side channels that have a width of 180 μm and a height of 100 μm. It also contains three channels in the middle, where two channels have a width of 400 μm and a height of 100 μm, and one channel that have a width of 120 μm and a height of 100 μm. A glass slide was used as a bottom part of the device to provide better image quality.

**Fig 1:**
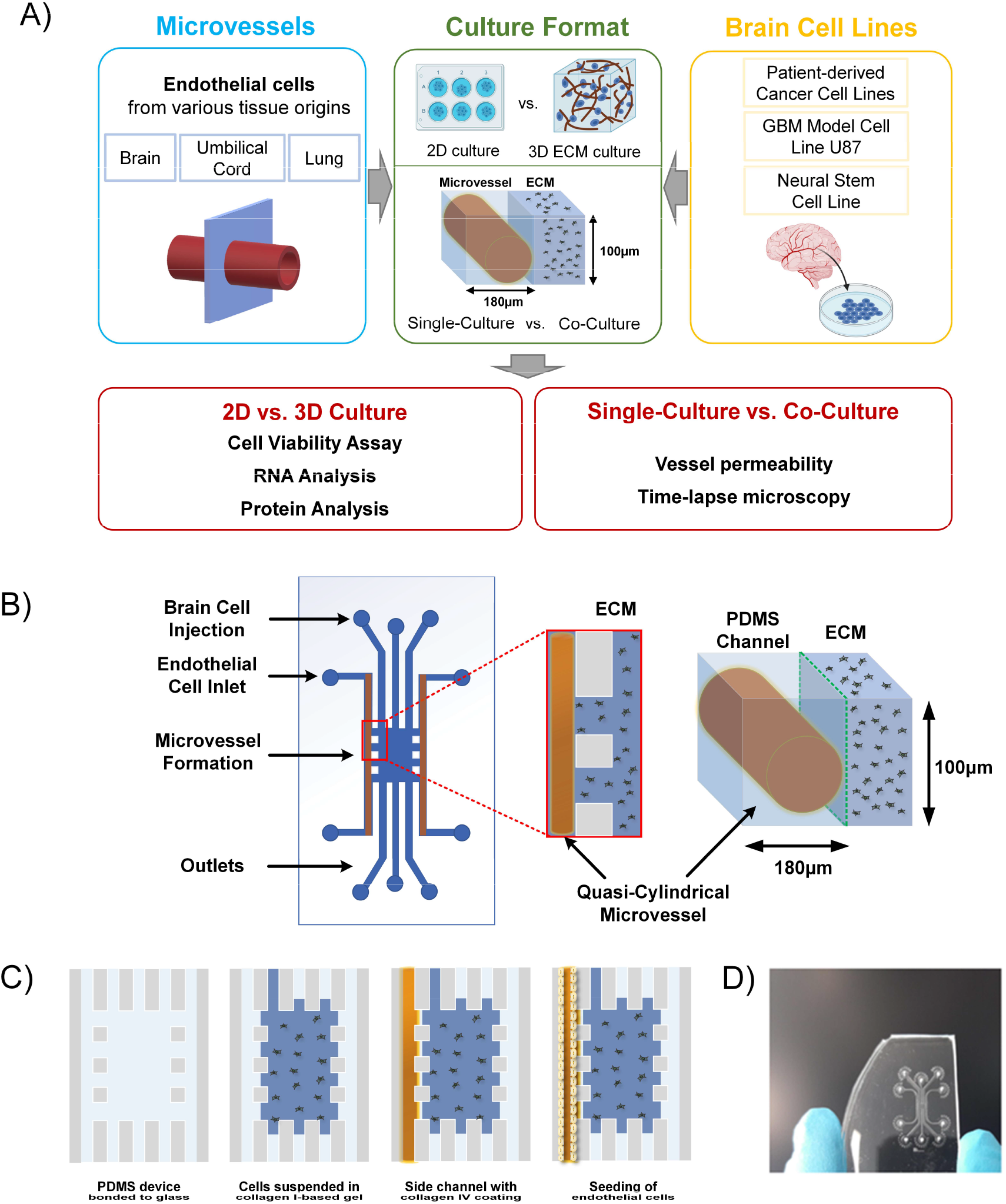
A) Schematics demonstrating physical attributes of on-chip perivascular niche with patient-derived glioma cells. B) Microfluidic device design and co-culture configuration. The design scheme of the microfluidic device, where the ECM-integrated PDMS channel shelters the quasi-cylindrical microvessel formed with various endothelial cells; C) cell suspension injection leads to two vacant side channels, and a central 3D ECM cell culture chamber, then seeding of endothelial cells as a side channel to form a microvessel; D) Photograph of the actual microfluidics channel fabricated for microvessel-on-a-chip implementation.

The microfluidic master was fabricated using the soft lithography process. SU-8 negative photoresist (MicroChem) was coated onto a 6” silicon wafer *via* spinning coating. Mask was then patterned using standard ultraviolet exposure. The microfluidics channels were fabricated by molding pre-crosslinked PDMS on the master. PDMS resin (Sylgard, 10:1 w/w ratio between elastomer and curing agent) was poured on the master and desiccated to remove the bubbles formed during the mixing process. The PDMS was then placed in an oven for 3 hr at 65 °C until fully cured. Afterwards, the PDMS was peeled off, and inlet and outlet holes with diameters of 0.75mm were punched. Foreign particles were removed from the PDMS surfaces using adhesive tape, washing with ethanol, and then blow-drying. The 3mm thick PDMS was bound to a round glass cover slip with a diameter of 22mm by air-plasma treatment (Femto Science; 15 sec, 25 sccm, 10 power), forming the microfluidic device.

### 2.2 Coating of microfluidic channels

Plasma treatment conferred hydrophilic properties to the channel walls, which made the subsequent poly-D-lysine (PDL) coating process feasible. The microfluidic channels were coated with PDL (1 mg/ml in deionized water (DIW)) for 5 hr at 37 °C. After coating, residual PDL was washed out multiple times using DIW to remove any excess molecules that could cause cellular damage. The device was then placed at 50 °C for 18 hr to recover the surface hydrophobicity. The hydrophobic recovery was necessary to successfully insert collagen I gel in the central region of the device, forming vertical barriers facing the cell culture channels.

### 2.3 Incorporation of an extracellular matrix

Collagen gel was incorporated in the PDMS-based device to establish a 3D extracellular matrix for cancer cell culture. While levels of collagens in the normal adult brain are relatively low, in glioma, collagen levels are elevated and play an important role in driving the tumour progression^20^. The ECM of brain tumours consists of the basement membrane components, collagen IV, laminin and fibronectin lining the blood vessels, as well as collagen I, tenascin-C, vitronectin and hyaluronan surrounding the tumour^20–22^. Thus, collagen I-based hydrogel and collagen IV coating for the outermost side channels were chosen for the microfluidic platform constructed.

### 2.4 Hydrogel injection and side channel coating

After restoring hydrophobicity in a PDL-coated device, the microfluidic chip was glued with a silicon rubber compound (RS Components, 692-542) to a new, bottomless polystyrene dish and collagen I gel was prepared and carefully injected from the access port of one of the channels directly facing the side channels. In terms of the gel preparation, DIW, collagen I rat protein (Gibco, A1048301), x10 phosphate-buffered saline (PBS; Gibco) with phenol red and 0.5N NaOH (Sigma-Aldrich) solutions were kept on ice. Using a wide orifice pipette tip, 27.2 μl of collagen was added to a 0.5ml tube kept on ice. 5 μl of x10 PBS with phenol red and 3 μl of 0.5N NaOH was added in a new 0.5ml tube and mixed. 5.4 μl of DIW was then added and mixed, until the gel appeared to be uniform in colour. Following the above procedure, the collagen gel concentration was 2 mg/ml and the pH around 7.5, resulting in a pink colour solution. Next, the gel solution was carefully injected into the access port of one of the channels directly facing the microvessel channels and interfaces were created between the cell culture channels and the central region of the device. Gel cross-linking was performed at room temperature for 1 hr. The gel was able to distribute in the central region and in the three channels connected to it, leaving the two side channels empty for endothelial cell seeding. Vertical interfaces could be created at the gaps between the pillars separating the side channels from the central region. 10 min after gel insertion, drops of cell medium were positioned on top of the access ports of all the channels, except for the outermost side channels. After gel cross-linking, collagen IV (Sigma, C5533, 1 mg/ml in DIW) was inserted inside the side channels to perform a second coating of the channel surfaces with a substrate suitable for cell seeding. Cell medium was added to the polystyrene dish, surrounding the device, to avoid solution evaporation inside the channels and the device was left inside an incubator, at 37 °C, for 1.5 hr. After collagen IV coating, the side channels were washed several times with cell medium to remove any excess of uncoated collagen. Next, the device was completely submerged in cell medium and stored inside the cell incubator for endothelial cell seeding.

### 2.5 Cell culture

#### 2.5.1 U87 – human GBM cell line

The U87 human GBM cell line was purchased from the American Type Culture Collection (ATCC). For maintaining the cell line, cells were cultured in T75 flasks in Dulbecco’s Modified Eagle’s Medium (DMEM, Gibco), supplemented with 10% foetal bovine serum (FBS, Gibco) and 1% penicillin/streptomycin (Sigma) at 37 °C, in a humidified 95% air and 5% CO2 atmosphere. They were split after reaching 80-90% confluency. Confluent T75 flasks were used for experiments. After trypsinization and neutralization with cell culture medium, cells were counted. Next, the spun down cell pellet was re-suspended in a volume of cell medium that brought around 500 cells per 5 μl. For this step, endothelial growth medium (EGM-2MV, Lonza) was used. Seeding density of 100,000 cells/ml was chosen as a compromise between mimicking the low number of tumour cells of the *in vivo* condition and increasing the chance to have at least one tumour cell to analyse in any given ROI. Meanwhile, the hydrogel was prepared via the same method as described in the previous section. 3 μl of cell suspension of hydrogel was then injected using one of the middle access ports. The device was then placed in an incubator and drops of cell medium were added 10 min later. 5 min after that, the device was flipped upside down to make sure the cells did not settle down on the bottom but suspended across the hydrogel layer. 30 min after hydrogel injection, collagen IV was perfused in the side channels. 1.5 hr after collagen IV coating the outermost side channels were washed, and the device was submerged in cell culture medium. In parallel, cells were cultured in 3D collage type I gel droplets in μ-slides (ibidi, IB-81506). μ-slides have small chambers where 5 μl of ECM-embedded cells can be cultured with 40 μl of medium to sustain the culture.

#### 2.5.2 Patient-derived human GBM cell lines

All patient-derived human GBM cell lines, were obtained under an MTA, from the Glioma Cellular Genetics Resource (www.gcgr.org.uk) funded by Cancer Research UK, and from the Pollard Lab at University of Edinburgh. The culture protocol for the GBM cell lines were detailed elsewhere ^23^. For the healthy brain control, the foetal neural stem (FNS) cell line GCGR-NS6FB_A was used. The FNS cell line was also obtained from the Glioma Cellular Genetics Resource.

| ***Glioma cell line name*** | ***Classification*** |
| --- | --- |
| *GCGR-E13* | classical |
| *GCGR-E17* | classical |
| *GCGR-E28* | classical |
| *GCGR-E35* | mesenchymal |
| *G166* | classical |

#### 2.5.3 Endothelial cell cultures

The hCMEC/D3 were commercially obtained (VH Bio) and cultured in endothelial growth medium in flasks coated with collagen from calf skin (Sigma, C8919; 5% v/v). Cells were cultured in standard culture flasks at 37 °C, in a humidified 95% air and 5% CO_2_, and the medium (EGM-2MV) was changed every other day until seeding. All experiments were conducted using hCMEC/D3 of passage between 27-30. Pooled HUVECs were purchased from Promocell. HUVECs growth medium was made up with endothelial basal medium, necessary growth factors and supplements (EGM-2MV). Cells were cultured in standard culture flasks coated with 5% v/v collagen from calf skin, at 37 °C, in a humidified 95% air and 5% CO_2_, and the medium was changed every other day until seeding. Cells were split typically once per week after dissociation with Trypsin solution and centrifugation. All experiments were conducted using HUVECs of passage 6 or lower. HMVEC-L were purchased from Lonza. The cells are a mixed population of both blood and lymphatic endothelial cells. HMVEC-L growth medium is made up with endothelial basal medium, necessary growth factors and supplements (EGM-2MV). Cells were cultured in standard culture flasks coated with 5% v/v collagen from calf skin, at 37 °C, in a humidified 95% air and 5% CO_2_, and the medium was changed every other day until seeding. Cells were split typically once per week after dissociation with Trypsin solution and centrifugation. All experiments were conducted using HMVECs-L of passage 6 or lower.

#### 2.5.4 Cell seeding for microvessel formation

A suspension at a density of 5×10^6^ cells/ml was mixed well to avoid cell clumping. Then, 3×l of the cell suspension was injected inside the cell culture channel by manually decreasing the pipette volume; therefore, the cell seeding was performed at a very low flow rate (~0.2 ×l/s), which was necessary to obtain a uniform distribution of cells inside the channel. It is to note that in order to avoid any back flow of cell solution inside the channels, the device had to be submerged in medium. The device was then stored inside the incubator for about 2 hr, to allow the cells to attach to the bottom surface of the channel. Next, a second seeding of endothelial cells was performed, and the device was turned upside down to allow the cells to attach to the top surface of the side channel. The device was put back to the upright position after 1 hr.

### 2.6 Vessel perfusion

Vessel perfusion was achieved through gravitational-driven flow by the following procedure. A 3D-printed stage was designed to fit a ∅100 mm polystyrene dish as a medium reservoir and two ∅35 mm with microfluidic devices, which were connected by tubing (Fig S2). Flow rates in the outermost channels were found experimentally and showed to be reproducible and reliable within the same devices and between devices. The results showed that a height difference of 4 mm could be used to achieve a flow rate of around 1.5 μl/min (Fig S2). Shear stresses were calculated using Computational Fluid Dynamics over a range of flow rates based on the results generated empirically. The results demonstrated that the flow velocities and shear stresses over the collagen interface were within the desired range. Shear stresses between 1.4 and 0.4 dyne/cm^3^ were calculated within the side channel of the device with velocities around 1-2 mm/sec (Fig S2).

### 2.7 Cell viability tests

For testing cell viability, LIVE/DEAD™ Viability/Cytotoxicity Kit (Life Technologies, L3224) was used. Briefly, 20 μl of 2 mM ethidium homodimer-1 (EthD-1; enters cells with damaged membranes) stock solution was mixed in 10ml of sterile PBS. Next, 5 μl of 4 mM calcein AM (CAM; live cells with ubiquitous intracellular esterase activity have the ability to enzymatically convert the virtually non-fluorescent CAM) stock solution was added to the 10 ml EthD-1 solution and mixed. That resulted in final concentration of 4 μM EthD-1 and 2 μM CAM. 200 μl of the prepared solution was then added on top of the microfluidic device and perfused through the lateral microchannels. After 25 min of incubation at room temperature, cells were visualized by confocal microscopy – viable cells (CAM-positive) staining green and dead cells (EthD-1-positive) staining red. Cell viability profiles were then evaluated by analysing the fluorescence intensity of the viable/dead cells across the microchambers, and calculating the ratio of live to dead cells. All confocal images were taken at different focal planes (2.5 μm step size) with subsequent image analysis performed using FIJI software.

### 2.8 RNA analysis

#### 2.8.1 Cell recovery from microfluidics devices

In this study, a protocol for endothelial cells extraction after they had been cultured in a microfluidic device (either mono- or co-culture) was optimized. Accutase® was used to detach endothelial cells in the outermost side channel. PicoPure™ RNA Isolation Kit (ThermoFisher, KIT0204) was used to isolate the RNA from cells extracted from microfluidic devices. First, all culture medium was removed from the dish with the microfluidic device of interest. Then 3 μl of Accutase® was added to the microvessel channel. After that, the dish containing the microfluidic device was placed inside the incubator for 1 min. After confirming that the cells have detached using a brightfield microscope, cells were removed from the channel and added to a 1.5 ml tube. 1ml of cell culture medium before spinning cells down, and then the protocols of PicoPure™ RNA Isolation Kit was followed. Once extracted as directed, RNA concentration and purity were assessed by absorbance measurements using NanoDrop 2000c (ThermoScientific). In every experiment, RNA from cells recovered from microfluidic devices and 2D control samples was checked, to make sure pure RNA was obtained. Next, to assess mRNA expression levels, 500 ng of total RNA was reverse transcribed (complementary DNA (cDNA) was synthesized using PrimeScript™ RT-PCR Kit (Takara) according to manufacturer’s instructions) and analysed by qPCR. Reactions for each sample were performed in triplicate using a PCR protocol (95 °C activation for 10 min, followed by 40 cycles of 95 °C for 15 s and 60 °C for 1 min, and 4 °C cooling hold step) in an ABI StepOnePlus Detection System (Applied Biosystems). ΔΔCt values for genes were examined using Ct values generated by StepOnePlus software (Applied Biosystems).

#### 2.8.2 Cell recovery from 2D cell culture

RNA from cells was isolated using TRIzol reagent. Typically, 6-well plates were used for the experiments, but the volumes were adjusted accordingly to different plate sizes. First, the medium was aspirated from the plate and cells were lysed using 1 ml of TRIzol reagent per well, which was incubated with the sample for 5 min at room temperature. Then, 200 μl of chloroform was added, and the sample was vortexed and incubated for 15 min at room temperature before centrifuging at 4 °C for 15 min at 12,000 rpm. 150 μl of the colourless top phase (containing DNA) was transferred to a new tube. Next, RNA was precipitated by adding 500 μl of isopropanol. The sample was vortexed and incubated at room temperature for 10 min, then centrifuged at 12,000 rpm for 10 min at 4 °C to obtain a pellet, which was then washed with 1ml of 75% ethanol. After centrifugation at 12,000 rpm for 3 min, the sample was air-dried for 5 min at room temperature. Finally, the pellet was dissolved in maximum 30 μl of RNase-free water.

#### 2.8.3 Cell recovery from 3D cell cultures

Collagenase P (Roche, 11213857001) was dissolved in PBS to a final concentration of 8 mg/ml. Collagenase solutions were subsequently sterile-filtered and added to the hydrogels. For degradation of hydrogels in μ-slides, 30 ×l were added per well. Collagenase incubation was performed at room temperature for 10 min. Afterwards, the recovered cell suspension was transferred to a fresh tube and washed with PBS before subsequent steps. PicoPure™ RNA Isolation Kit was used for RNA isolation, as per manufacturer’s recommendations. Then, RNA concentration and purity were assessed by measuring 260/280 absorbance ratio. Pure RNA produces 260/280 ratios around a value of 2, and lower values are indicative of high protein content in the sample. For an example experiment, the average ± SEM 260/280 ratio values for 2D and microfluidic system were 1.952±0.025 and 1.949±0.037, respectively. In each experiment, results obtained from samples cultured in 3D ECM, as compared to 2D setup, passed Shapiro-Wilk normality tests with showing no significant differences in quality of obtained RNA.

#### 2.8.4 Reverse transcription PCR

The RNA samples had to be reverse transcribed to obtain cDNA from isolated RNA. PrimeScript™ RT-PCR Kit was used for cDNA synthesis. The reaction was set in a 0.2 ml PCR tube by using 4 μl 5x cDNA synthesis kit buffer, 1 μl PrimeScript™ enzyme mixture, x μl nuclease-free water, and y μl 500 ng RNA sample. Next, the following thermal cycle was run: 5 min at 25 °C, 30 min at 42 °C, 5 min at 85 °C, hold at 4 °C. Afterwards, 30 μl of RNase-free water was added to all samples. The aliquots were frozen at −20 °C for short, and −80 °C for long term storage.

#### 2.8.5 Quantitive PCR

In order to assess differences in expression of molecules of interest, quantitative polymerase chain reaction was carried out. SYBR® Green was used as a fluorophore. SYBR® Green is a fluorescent cyanine dye that has high affinity for double-stranded DNA. The reactions were set up in 96-well PCR plates under semi-sterile conditions (laminar flow) in order to avoid contamination. The reaction was set up by adding 5 μl SYBR® Select Master Mix, 0.75/075 μl fw/rv primer (10 μM), 2.5 μl ddH2O, and 1 μl cDNA. Obtained cycle over threshold (Ct)-values were analysed by normalizing them against a housekeeping gene (*GAPDH*), and then comparing the differences with a control, which was either set or chosen based on the lowest relative expression (ΔΔCt-method).

### 2.9 Protein analysis

#### 2.9.1 Western blotting

For protein analysis *via* western blotting, cells were seeded either inside microfluidic devices, on 6-well plates, or μ-slides (3D ECM culture). To harvest cells from a microfluidic chip, cells were detached and collected in a tube, then spun down, and a cell pellet was subjected to protein analysis after cell lysis in 50 μl of RIPA buffer with phosphatase and protease inhibitors. To extract cells from collagen I-based hydrogel, collagenase was used and then the obtained cell pellet was subjected to protein analysis after cell lysis in 50 μl of RIPA buffer with phosphatase and protease inhibitors. For a 6-well plate, the plate was first chilled on ice, then the medium was removed from the plate and the cells were washed twice with chilled PBS. 300 μl of RIPA buffer was then added to each well and incubated on ice for 5 min to allow cells to lyse. The next steps were the same for both cells cultured on a 6-well plate, microfluidic devices and μ-slides. Thus, the solution was collected into 1.5 ml tubes, which were centrifuged at 14,000 rpm at 4 °C for 10 min and the supernatant was either used immediately or frozen and stored at −80 °C. Bio-Rad protein assay was used to measure protein concentration against a BSA standard. Then, 6x Laemmli sample buffer was added, before boiling the samples for 2 min at 95 °C. This was done to denature the protein. Prepared samples were loaded onto polyacrylamide gels. Next, SDS gel electrophoresis (SDS-PAGE) was used to resolve the samples, which were then transferred to polyvinylidene fluoride (PVDF) membranes (Millipore). After the transfer, membranes were blocked for 1 hr at room temperature in 5% milk powder in TBS/T (1M Tri-HCl, 5M NaCl, 1% Tween-20, DIW). Next, the primary antibody was diluted in 1% BSA (Bovine Serum Albumin) in TBS/T and membranes were incubated in this solution overnight at 4 °C, before being washed 3 times in TBS/T. For each wash, the membranes were placed on a shaker for 5 min. The secondary antibody was diluted in TBS/T and incubated with the membrane for 1 hr, and then washed twice. The antibody was detected using the ECL plus reagents (GE Healthcare). Detection was done in a dark room, and exposure time varied according to intensity of signal. Where indicated, densitometry was acquired using ImageQuant TL software.

#### 2.9.2 Immunofluorescence

For analysis of protein expression *via* immunofluorescence, cells were fixed and stained with fluorescent antibodies. We used immunofluorescence for staining cells grown in: i) 6-well plates, ii) ibidi μ-slides, or iii) microfluidic devices. In all cases, cells were fixed in 4% paraformaldehyde for 20 min at room temperature, washed 3 times in PBS and then permeabilized with 0.2% Triton™ X-100 (Sigma) in PBS for 3 min. Cells were washed again 3 times in PBS, and then blocked by incubating the cells with 4% BSA in PBS for 1 hr. Next, cells were incubated with primary antibodies, as listed in the Table S1 A), at the appropriate dilution overnight at 4 °C. The next day, cells were washed 3 times with PBS and then incubated with a secondary antibody of choice, as listed in Table S1 B), as well as nuclear and/or F-actin counterstain for 1 hr at room temperature in the dark. Cells were washed 3 times in PBS before imaging with a Leica SP5 confocal microscope was performed.

For experimental sets where live/dead cells were being studies, the images were analysed by quantifying labelled cells and by measuring the intensity of fluorescence. The analysis was carried out *via* FIJI. To quantify the number of marker-positive cells, the plugin *cell counter* was used after gating out weakly stained cells with the threshold. In order to quantify fluorescence intensity single cells were outlined and measured with the *measure* tool, which displayed the values for area, mean, internal density and raw internal density. As a control, sections of the background were outlined as well. In order to calculate fluorescence, the following formula was utilised:

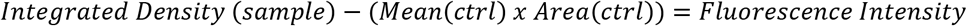

For fluorescence intensity and quantification, images were taken with the same settings (exposure, saturation, intensity) for each label and magnification in order to accurately compare them. Moreover, different cell size was taken into account for the analysis.

### 2.10 Live-cell imaging and image analysis

An in-house image analysis programme to pre-process the time-lapse images and create a database to automatically classify cell interactions with the regional microenvironments was used. This novel program utilizes several open-source applications, such as FIJI and CellProfiler (FIJI: https://imagej.net/Fiji; Cell Profiler: https://cellprofiler.org/), and merges them with focus stack (z-stack) image analysis to enable cell tracking and segmentation. The experiments used brightfield (or transmission) images and green fluorescence protein (GFP) images. Intensity of GFP expression of the z-stack images acquired during the time-lapse experiments was projected onto one x-y plane at each time point; thus, the projected cell area, that is the 2D projection of the 3D cell shape, was obtained at each time point. Simultaneously, brightfield images were used for correcting the cell outline, allowing the programme to recognize individual cell bodies. Afterwards, the ROI manager plugin of FIJI, applied to the brightfield images of the z-stacks, was used to define ROIs – gel and interface between 3D ECM cancer cell culture and microvessel – within each field of view. The Manual Tracking plugin of FIJI, applied to the sequence of images, was used to track the position of cancer cell nuclei over time. Targeting the cell nucleus while tracking cell movements is crucial, to enable the image analysis software (CellProfiler) to perform the segmentation of the cancer cell. During segmentation, the cell shape is separated and differentiated from the background. Segmentation allowed to extract the cell shape features in the previously defined regions at each time point. Furthermore, the z-coordinate of each cancer cell was calculated by finding the z-level of the maximum GFP signal within the projected cell area. Cell segmentation combined with maximum GFP analysis determined the cell centre. All the obtained information was used to describe cell behaviour in the presented system. The final segmentation presents an error of about 15% on the projected cell area of each cell and an error of about 2 ×m (in both x and y direction) in the location of the cell nucleus.

Cell speed, velocity and polarization were measured and analysed using the image-assisted microfluidic platform. The parameters were calculated for 3D cancer cell culture in the presence of a microvessel, or in a reference system – without endothelial lining. Cancer cell migration velocity (measured between two consecutive time points) was quantified. The direction of the cancer cell migration was quantified by velocity components in the x-direction (perpendicular to the endothelium-collagen matrix barrier) and y-direction (along the channel direction).

The presence of asymmetric shape features in cancer cells with respect to the position of the cell nucleus was described by using the cell polarization parameter. The position of the cell centre of mass was calculated by the CellProfiler software, weighting the x and y coordinates of the pixels within the projected cell shape with their GFP intensity; a computational error of about 20% was made due to misrecognition of the projected cell shape profile during automatic segmentation.

### 2.11 Vessel permeability measurement by dextran diffusion

Upon formation of an endothelial monolayer, the diffusive permeability was measured with fluorescently labelled dextran (fluorescein isothiocyanate-dextran, FITC-dextran) in culture medium. The permeability measurements and calculations were adapted from Funamoto *et al*. ^24^ Briefly, an intact endothelial monolayer gave rise to an intensity drop between the channel and the gel region once the fluorescently labelled dextran was introduced and persisted over time as dextran slowly diffused across the monolayer into the gel. Hence, the microvessel permeability was assessed by observing the diffusion of 70 kDa FITC-dextran (Sigma, 46945; 7 mg/ml in PBS, diluted further 1:9 in cell culture medium) from the microvessel channel into the collagen gel. The diffusion experiments were performed in the environmentally controlled chamber (set at 37 °C, 5% CO_2_) of a fluorescent microscope (Leica, CTR6500). The dextran solution was perfused inside the microvessel channel and images were acquired every 30 sec over 30 min. Beyond 30 min, the geometry of the device (e.g., pillars and the limited size of the gel chamber) broke down the *perfect sink* condition assumed in the calculation. Obtained fluorescent images were analysed using an open-source software (FIJI).

### 2.12 Statistical analyses

Data were analysed using GraphPad Prism software and its built-in tests, or RStudio. Data were tested for normality with Shapiro-Wilk test and subsequently analysed with suitable parametric or non-parametric tests. In order to assess significance, unpaired or paired t-test or ANOVA with Tukey’s post-hoc test was used, depending on the experiment in question. The following asterisk rating system for p-value was used: *p≤0.05, **p≤0.01.

## 3 Results and Discussions

### Protocol optimization to study gene and protein expression in cells cultured in 3D microfluidics

We fabricated an *on-chip* vascular niche (Fig 1), to investigate the interactions between patient-derived glioma cell lines and different organ-specific endothelium. A single, quasi-circular perfusable microvessel is cultured adjacent to a 3D ECM cancer cell culture, which adds the vessel architecture, shear stress, and ECM components compared to commonly used co-culture set up like Transwell® systems. We optimized the fabrication pipeline in order to prepare a batch of devices (typically 25 devices) ready for cell culture in 3 days. Despite the advantages of cell culture in microfluidic devices, that would better emulate *in-vivo* conditions than conventional 2D cell cultures used to study cancer cell interactions with vascular microenvironment, the integration of these techniques as a research tool in the mainstream biology research has been slow^25^. Most of the microfluidic devices available on the market offer a low number of read-outs in assays, which are generally derived by microscopy analysis. Genomic or proteomic analysis post microfluidic cell culture require protocol optimisation to retrieve cells in 3D culture from the microfluidic device. Here, we optimized several protocols for biological analysis on cells post-3D culture in microfluid device, as summarized in Fig 1, which make our platform flexible and adapt to be employed in the most used biological analysis, such as protein/gene expression analysis coupled with functional analysis such as vessel permeability and live-cell imaging of cancer cell migration.

The study of the glioma-associated neovascularization mechanisms have been an ongoing research topic since microvascular proliferation was declared a pathological hallmark of malignant high-grade gliomas ^26,27^. We developed a quick and straightforward method to recover endothelial cells from our *on-chip* vascular niche coupled to microfluidic assay, to perform qPCR analysis. After cell isolation, we used a commercial RNA isolation kit, RNA concentration and purity were assessed by absorbance measurement (260/280 ratio). RNA from cells recovered from microfluidic devices and 2D control samples was checked to ensure pure RNA from cells was obtained. The results passed Shapiro-Wilk normality test and were then subjected to a *t-*test. Our analysis between the 2D and 3D endothelial cell RNA quality test shows no significant difference, indicating that our RNA extraction and purification protocol is comparable to 2D assay routinely used in research. Using our optimized method, it was possible to extract up to 75 ng/μl of RNA purity within acceptable range. The extracted cells from the 3D microfluidic device are pure endothelial cell population as confirmed by qPCR analysis. It has shown that they did not contain any GFAP, a particular marker for the GBM cells.

### 3.1 Brain cancer cells in 3D ECM and microvessel-on-a-chip configuration

#### Glioblastoma U87 cell culture in 3D collagen-I gel

The U87 cells are commonly used to investigate GBM, therefore here we optimised the protocol to grow them in 3D collagen I-based ECM cell culture. In previous *in vivo* studies, it has been reported that U87 cell type failed to accurately model human GBM compared to patient-derived tumour stem cells. They described that U87 cells, cultured in 2D *in vitro* cell culture with FBS-enriched medium, exhibits low levels of neural stem cell markers, such as nestin, SOX2, and CD133 ^28,29^. Here, we set up a protocol to culture the U87 cells in 3D-ECM with important components of the microenvironment characteristic of glioblastoma stem-like cells ^30,31^, such as EGF, FGF-2 and laminin in serum-free medium.

To optimize the 3D ECM cell culture conditions for U87 cells, we compared cells grown in collagen I-based hydrogel embedded in μ-slides maintained either with 10% FBS/DMEM or serum-free medium. The U87 cells cultured in either 10% FBS/DMEM and EGM-2MV media behaved similarly, indicating that the culture media does not impact the cell growth in 3D culture conditions (Fig S3). 3D culture of U87 cells after 3 days showed that around 90% of the cells were viable (Fig S4 A) & B)). Furthermore, we compared if there were differences between the 2D and 3D ECM culture systems, in serum-containing versus serum-free media conditions at mRNA level (Fig S4 C) and D)) by seeding the same number of cells in either 2D (flat, glass bottom) 96-well plates, or in collagen I-based hydrogel cultured in μ-slides. No differences in gene expression of cells grown in 10% serum and serum-free medium was observed (Fig S3). Of note, the 3D ECM culture significantly enhanced expression of stemness-related genes compared to the 2D culture, which is consistent with data previously published regarding GBM-derived cell lines cultured in collagen gels ^32–34^.

#### 3D collagen-I gel culture preserves glioma stem cell-like phenotype in patient-derived cell lines

Although cancer cell lines are the standard models in the fields of molecular cancer biology and drug discovery ^35^, there have been concerns regarding their inadequate representation of the *in vivo* cancer cell behaviour ^35–37^. Considering the complexity of both inter- and intra-tumour cellular, genetic and molecular heterogeneity of GBMs ^38,39^, it is of interest to design a human model in vitro, that can be easily used for culturing different patient-derived cancer cell lines, and to dissect whether glioblastoma sub-types influence the process of neovascularisation. We used five patient-derived cell lines, containing stem-like cell population (G166, GCGR-E13, GCGR-E17, GCGR-E28 and GCGR-E35). The FNS cell line GCGR-NS6FB_A was used as a healthy brain control and the U87 cell line we used to set up the 3D ECM model, was used as technical control. Since these cell lines are standardly cultured in 2D *in vitro* conditions, it was crucial to test whether our 3D ECM setup would influence the cell properties of these five GBM patient-derived cell lines. Therefore, we used the collagen I-based hydrogel as the medium to compare the brain-derived cells cultured in 2D or 3D ECM environment. All the tested cell populations were successfully cultured in the 3D ECM setup for 3 days with the expected phenotype (cell morphology) (Fig 2 A) (left panels)). Relative gene expression (mRNA level) analysis was performed to study the differences between 2D and 3D culture in proliferation, stemness and differentiation. We did not detect the glial fibrillary acidic protein (GFAP), a marker of astrocytic differentiation in the G166, GCGR-E35 and FNS cell lines both in 2D and 3D cultures ^40,41^, by qPCR. In all cell lines, proliferation (Ki67 marker^42^) in 3D ECM culture was slightly decreased when compared to the 2D one. On the other hand, in 3D ECM cell culture the expression of stemness-related markers (CD44, CD133, nestin, olig2, sox2) was increased compared to 2D culture in all GBM patient derived cell types studied, however we did not observe striking changes of GFAP expression between the selected cell lines with GFAP expressed (Fig 2 A)).

**Fig 2:**
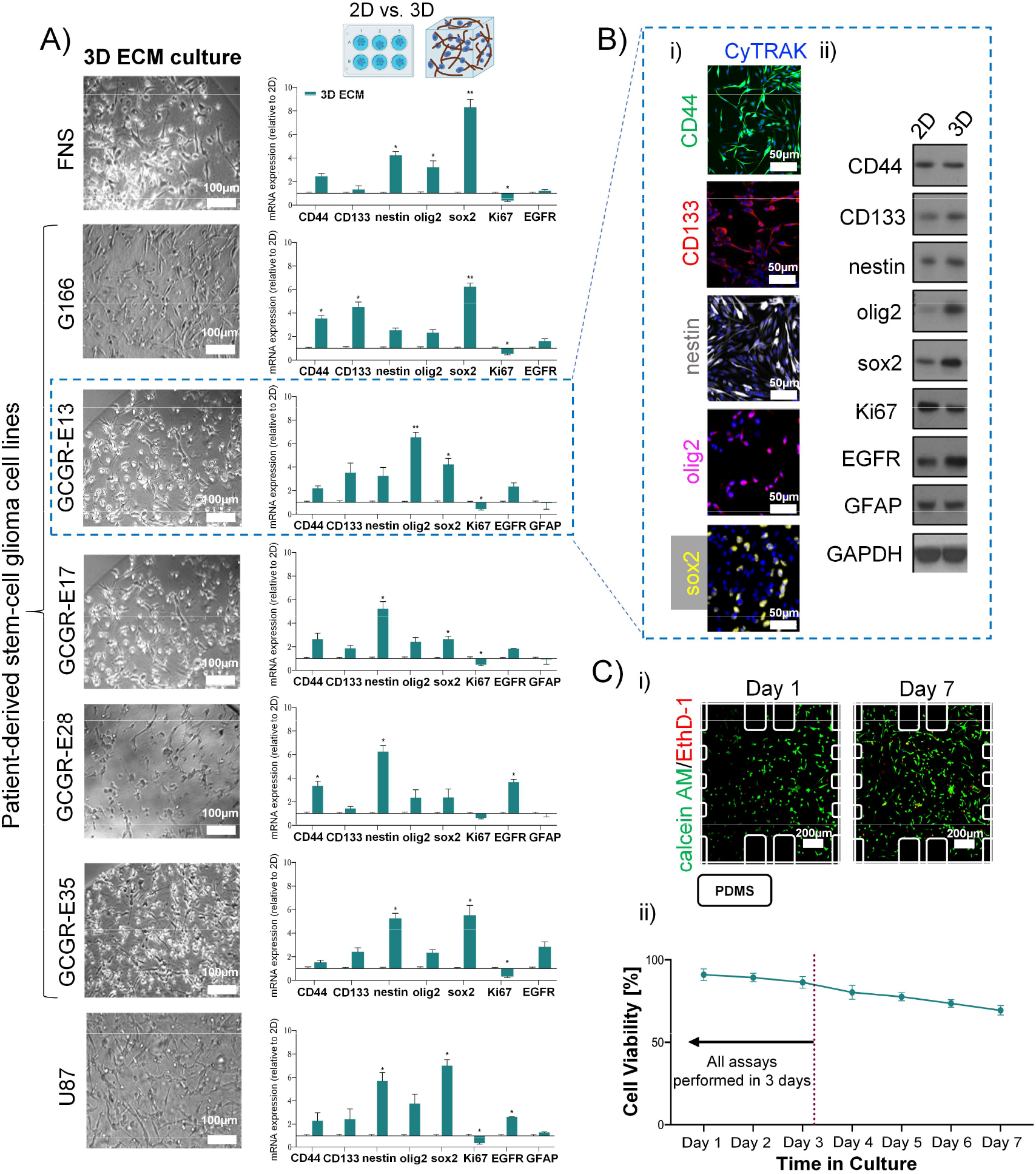
Patient-derived stem cell lines are characterised by stemness-related genes whilst cultured in 3D collagen I-based ECM for three days. A) Representative bright-field images of each GBM cell type selected cultured in 3D ECM acquired by light microscopy. mRNA expression of stemness-related gene in GBM cells normalised to 2D samples. Results for the 3D samples obtained from three independent experimental runs are presented by mean±SEM. Two-tailed t-test was used for significance. Note that for FNS, G166 and GCGR-E35, GFAP was not detected by qPCR; B) i) Example immunostaining images of GCGR-E13 cells grown in 3D for stemness-related markers: CD44, CD133, nestin, olig2, sox2. All were counter-stained for nucleus (CyTRAK); ii) Example western blot analysis for GCGR-E13 cells in relation to GAPDH, cultured in both 2D and 3D ECM; C) The viability of GBM cells in 3D culture. i) Example images of calcein AM/EthD-1 staining of GCGR-E13 cells at day-1 and day-7 culture, inside a microfluidic device; ii) Graph presenting the cellular viability over seven days in cell culture. Data presented mean±SD (percentage of live cells).

The stemness-related markers that we selected have been indicated for their role in maintenance properties of cell renewal and multi-lineage differentiation, contributing to phenotypic plasticity of GBM cells ^43,44^. Furthermore, epidermal growth factor receptor (EGFR) expression was upregulated in selected patient-derived cell lines (i.e. GCGR-E13, GCGR-E28). Overall, our mRNA level data are consistent with previously published data in GBM-derived cell lines derived from other sources, cultured in collagen gels ^32–34^.

We further confirmed the trend in differences between 2D and 3D ECM culture by western blot for a selected GBM cell line, the GCGR-E13 (Fig 2 B)), with the exception of CD44. CD44 mRNA levels were slightly upregulated at mRNA level, but not at protein level. Finally, we confirm the localization and protein expression of CD44, CD133, nestin, olig2 and sox2 in GCGR-E13 cells cultured in 3D ECM by immunostaining (Fig 2 B)).

The multi-level analysis of factors contributing to GBM phenotype has shown comparable trends in detection of mRNA transcripts and protein between 2D and 3D ECM cell culture. Based on the above results, we suggest the 3D collagen-I culture condition created for GSCs are suitable for maintaining their phenotype for 3 days.

#### Viability of GSCs in 3D ECM-integrated microfluidic culture

Next, we evaluated the cellular viability in the 3D ECM-integrated microfluidic culture. Each cell line was cultured within microfluidic devices for up to 7 days. Cells were cultured in the serum-free medium, and in the presence of gravity-driven flow. 21 devices (3 devices per day) for each cell line in triplicate (n=3, different days) were analysed. Cell viability of each cell line slightly decreased over time to over 70% of live cells after 7 days of culture. Two out of the seven cell lines maintained 80% of viability after 7 days of culture. Nonetheless, most dead cells at day 7, were found as part of necrotic clusters located near the PDMS pillars, at the centre of a device (Fig S5 B)). This could be due to a reduction of lack of nutrients and oxygen in that areas of the device, and a gradual increase of toxins accumulation inside the device. However, this area of the device is outside the region of interest to study the interaction between cancer stem like cells with cells forming 3D microvessels. Moreover, the experiments to study the cooperation between the brain cancer cells and the artificial microvessels did not require a culture longer than 3 days, time where all the cell types studied did not show a decrease in viability higher than 15%. Therefore, we can conclude that this microfluidic setup coupled with gravity-driven flow can support the 3D ECM culture of all five GBM patient-derived cell lines with acceptable level of viability.

### Microvessel-on-a-chip of different organ-specific endothelial cell types

#### Biological and functional characteristics of the microvessels

With the microfluidic set up, we next want to establish whether different types of organ-specific endothelial cells could influence the interactions between brain cancer cells and endothelial cells. With our method that allow rapid vessel formation, we investigated the human brain (hCMEC/D3), primary cell from umbilical cord (HUVECs) or lung (HMVEC-L) endothelial cells vessel formation in the 3D microfluidic device. With such capability of our device, we first asked whether there are differences between culturing endothelial cells as a monolayer in 2D versus in a 3D microvessel configuration. Each endothelial cell type was cultured on a collagen IV coated glass-bottom 6-well plates or in the 3D ECM-integrated microfluidic device. As control, we cultured endothelial cells on a standard polystyrene 6-well plates without collagen IV coating. Endothelial specific marker (VE-cad, CD31 and ZO-1) mRNA levels were measured (Fig 3 A)). Endothelial cells cultured on collagen IV coated substrate showed to have higher relative levels of endothelial markers, in particular the brain one (hCMEC/D3), as expected, however no significant difference between 2D and 3D culture was observed.

**Fig 3:**
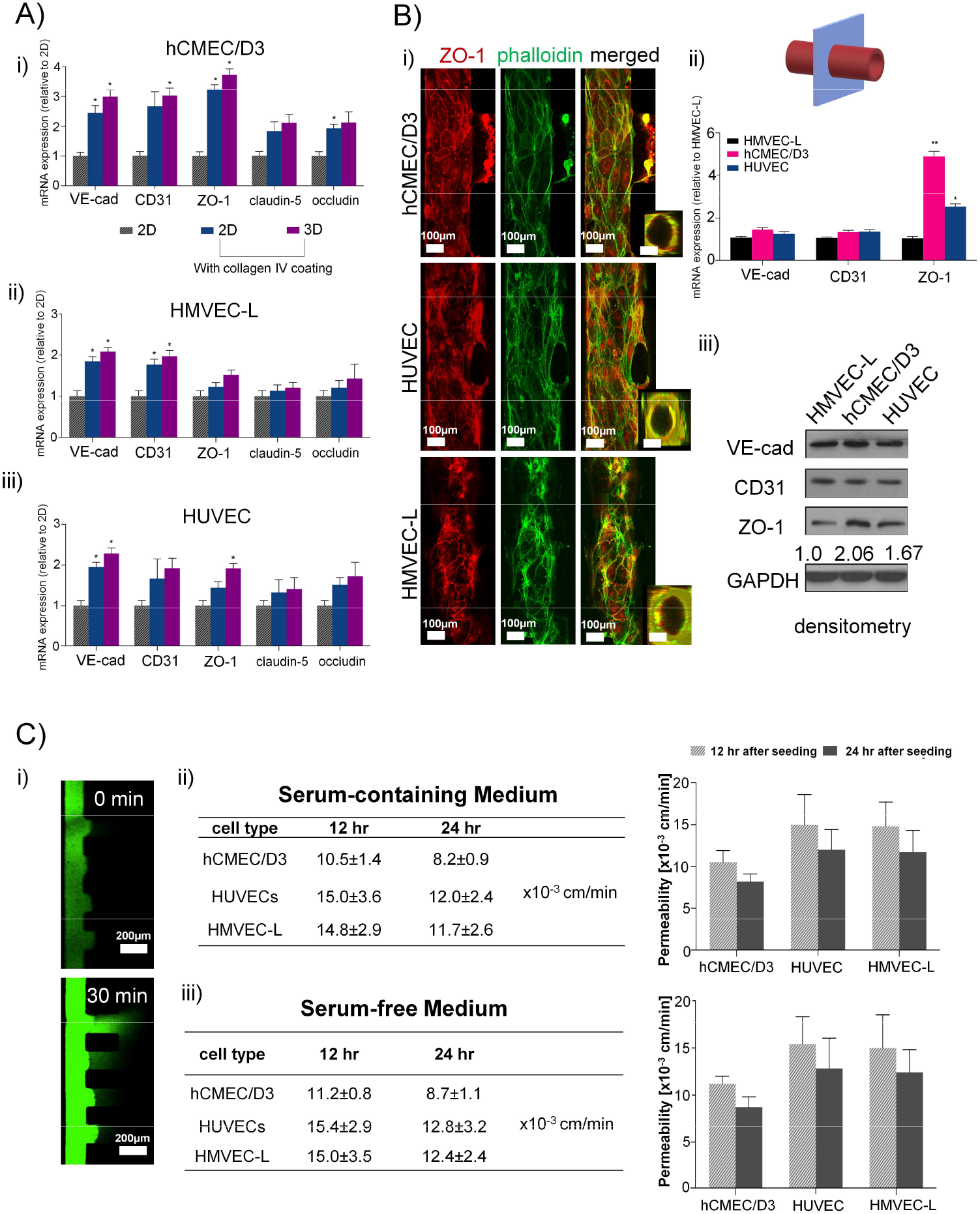
A) comparison of tightness-related gene expression in each endothelial cell type in 2D and 3D. mRNA expression normalised to 2D (without collagen IV coating) samples; results obtained from three independent experimental runs are presented by mean±SEM. One-way ANOVA with Tukey’s post-hoc test was used for significance; B) i) Microvessel lumen formed by various types of microvessels in contact with collagen I gel. Staining of junction protein ZO-1, cytoskeleton by phalloidin, and merged image. In all panels, on the right, a 3D reconstruction of the microvessel cross-section at a location along the vessel lumen (y-axis); ii) hCMEC/D3 express higher levels of endothelial tightness-related genes then HUVECs and HMVECs-L in the microfluidic setup. mRNA expression normalised to HMVEC-L (results obtained from three independent experimental runs are presented by mean±SEM. One-way ANOVA with Tukey’s post-hoc test was used for significance.); iii) western blot analysis in relation to GAPDH. Densitometry was acquired using ImageQuant TL software; C) Diffusion measurements showed appropriate level of permeability for the hCMEC/D3 vessel; i) Example images from permeability experiment showing diffusion after 30 min from 70kDa FITC-dextran injection; ii) Endothelial permeability measurements for cells in serum-containing medium 12 and 24 hours after seeding; and iii) Endothelial permeability measurements for cells in serum-free medium 12 and 24 hours after seeding.

As the concept of vasculature, in simplistic terms, it can be compared to a pipe. Therefore, we aimed to build a platform with a *quasi*-circular microvessel to mimic in vivo microvessel geometry that can be visualized easily, and that is able to form in very close proximity to the ECM-embedded brain cancer cells to create an environment more similar to an in vivo scenario than a Transwell® system for instance.

Next, we tested the ability of the three endothelial cell types to form a microvessel structure of quasi-circular cross-section under our culture condition and compared the properties of microvessels formed, in 24 hrs of microfluidic culture. Endothelial tight junction ZO-1 and actin (phalloidin) staining revealed that all three cell types were able to form a microvessel structure with a quasi-circular cross-section under 3D culture condition by immunofluorescence (Fig 3 B)). Using our 3D ECM device, HUVEC and hCMEC/D3 cells formed more defined circular microvessel structure compared to HMVEC-L cells. Then, we isolated the endothelial cells from the 3D ECM microvessel and we measured RNA relative levels and protein expression of the endothelial junction VE-cadherin (VE-cad) and of the endothelial marker CD31 in all three cell types (HUVEC, hCMEC/D3 and HMVEC-L) (Fig 3 B) ii)). Both qPCR and western blotting revealed that hCMEC/D3 is characterized by significant higher levels of ZO-1 expression than the other two cell types (Fig 3 B) ii)). This is expected as ZO-1 is required for tight junction formation ^45–47^ in brain endothelial cells forming the blood brain barrier, as they have the function to tightly regulate it. Therefore, we further investigated other molecules forming the tight junctions, claudin-5 and occluding in hCMEC/D3 ^48^. Indeed, claudin-5 and occluding transcript levels were higher in hCMEC/D3 than in HUVEC or HMVEC-L (Fig S6 A)).

As the process of neovascularization is important in the context of GBM, we compared the RNA levels of neovascularisation-related genes across the three cell types forming circular microvessels in 3D, and no significant differences in mRNA relative levels were detected (Fig S6 B)).

The permeability of the barrier in the three type of microvessels was measured by fluorescent dextran perfusion. Leakage was not observed in any of the three microvessel types. The permeability values showed that the brain endothelial cell line, hCMEC/D3, form the tighter barrier compared to HUVECs or HMVECs, and the permeability on the microvessel increases after 24h of culture in the 3D device. These findings further confirmed that the hCMEC/D3 cells form a more favourable microvessel than HUVECs or HMVECs-L to study brain vasculature and GBM cells as they are specific to brain, and that they are adequate to be used in the specific setup we described here. Nonetheless, all three cell types were able to create vessels of acceptable permeability rates, as compared to other microvessel-on-a-chip models where hCMEC/D3, HUVECs or HMVECs were used ^49, 50, 51, 52^. Overall, the brain endothelial cells hCMEC/D3 of brain origin are the more suitable cell type to form the microvessel, compared to HUVECs and HMVECs-L, as confirmed through hCMEC/D3 cells’ lower permeability coefficient, higher expression of adherens and tight junction-associated genes. Our results are in line with a recent study which compared *on-chip* microvessels made up of HUVECs with brain microvascular endothelial cell microvessel, suggesting that the source and specificity of endothelial cells should be taken into consideration whilst designing an *in vitro* model for organ-specific cell culture ^53^.

#### Serum-free environment does not influence permeability of microvessels

In microfluidic co-culture setups, it is important to find a type of medium that can be used for all cell types within the device, providing optimum culture conditions. Due to the known drawbacks of serum, for instance its unknown composition and very high batch-to-batch variability, there is a growing trend to replace it with a precisely defined alternative ^54^, which makes it easier to manipulate by controlling certain factors. Moreover, as mentioned before, stem-*like* cancer cells are typically cultured in serum-free medium ^55^, thus it was vital to test whether the endothelial cells retain their properties under a serum-free media condition.

We evaluated the impact of serum-free medium on the vessel permeability determined using the medium EGM-2MV for hCMEC/D3 and HMVEC-L cells and EGM-2 for HUVEC, which are routinely used for these specific endothelial cell culture. The permeability of the three different endothelial cell types in microvessel does not change when cultured in standard serum or serum-free medium L (Fig 3 C) iii)) confirming existing reports that microvessels cultured in serum-free conditions can exhibit adequate vessel barrier functions^56, 57, 58^.

### Organ-specific cancer cell-endothelium interactions

Although angiogenesis has been regarded as essential for tumour growth and progression, recent studies suggest that a variety of tumours can advance without angiogenesis, and their microcirculation may be provided by nonsprouting vessels^5^. Establishing an *in vitro* model that allows the study of interactions between circulating and or disperse cancer cells and nonsprouting microvessels, can be of value towards understanding mechanisms contributing to brain tumour progression. In the current work, we aimed to evaluate gene expression pattern changes in the microvessels of different tissue origin upon interaction with brain cells in the system we set and characterized (Fig 1–3). Therefore, three separate co-culture setups were established to study the interactions between either fetal stem cells or GSC cells with vessels formed by hCMEC/D3, HUVEC or HMVEC-L, in serum-free medium and gravity-driven flow.

#### Characterization of endothelial cell microvessel-GSC co-culture in the microfluidic system

We first evaluated the viability of various brain-derived cells co-cultured with the hCMEC/D3, HUVEC or HMVEC formed microvessel. Cell viability for all the different combinations of the microvessel-GSC co-cultures was between 95% and 75% after 3 days of co-culture (Fig 4 A) and B)) indicating acceptable cell culture conditions.

**Fig 4:**
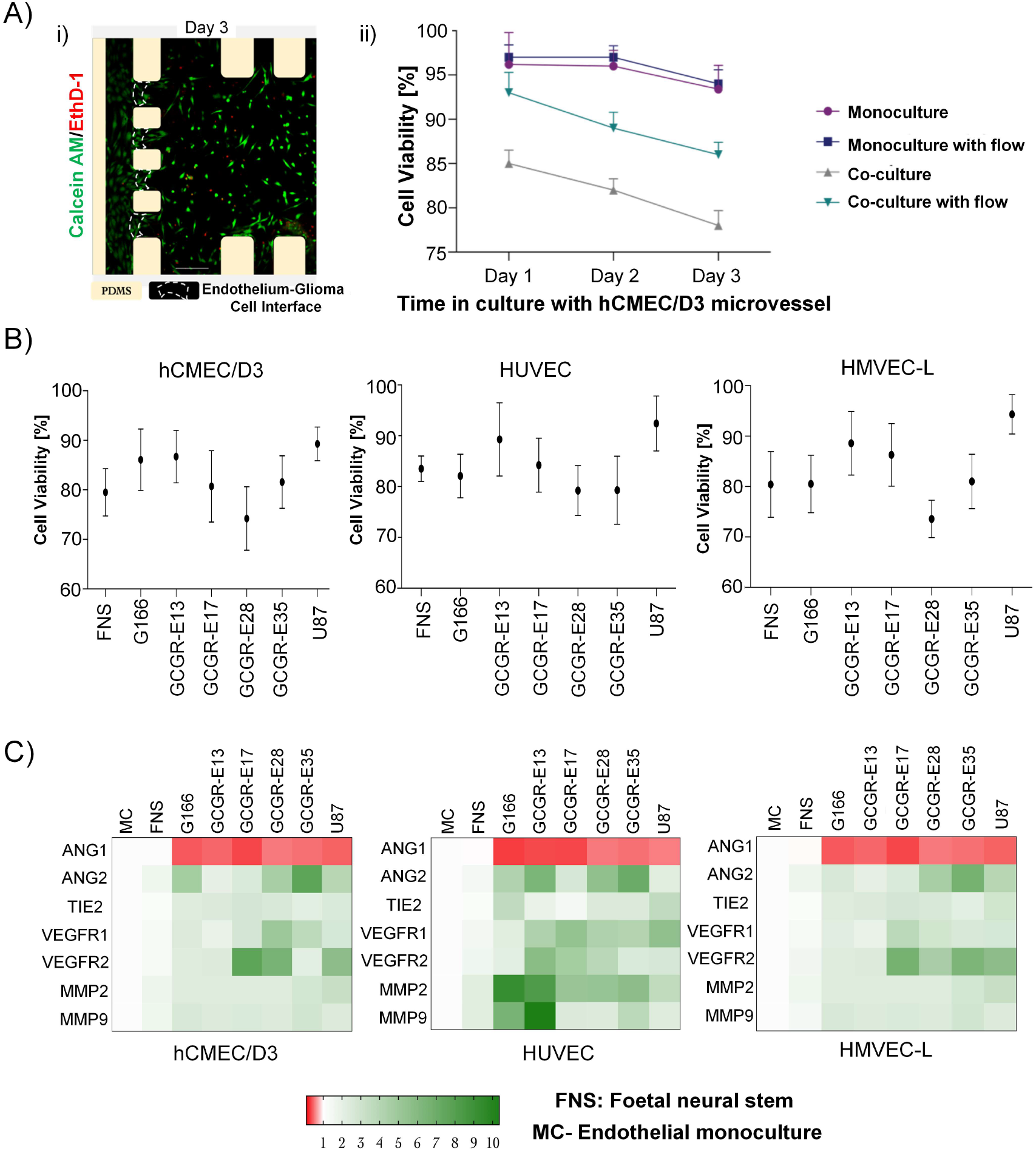
Viability of GSCs in 3D microfluidic device culture, with or without the presence of microvessels. A i) Example image of calcein AM/EthD-1 staining of GCGR-E13 cells at day three in co-culture inside a microfluidic device with gravity-driven flow; ii) Three-day GSC cell viability for four different configurations; B) Graphs presenting viability of six different GBM cell lines, a foetal neural stem cell line, and the U87 cell line at day-3 culture with different microvessel types. C) GBM cells induce different response patterns in endothelial cells in comparison to normal brain cells. Heatmaps were plotted in GraphPad Prism software after the analysis of mRNA expression in relation to GAPDH. Reference pattern was established from RNA isolated from endothelial cells cultured without the presence of brain cancer cells in microfluidic devices (MC-monoculture microvessel). Results obtained from three independent experimental runs. Green indicates upregulated expression, and red indicates downregulated expression.

Next, to determine how GSCs influence the microvessels, endothelial cells were recovered from microfluidic devices after 24 hrs of co-culture, to establish an expression pattern of neovascularization-related genes at mRNA level (Fig 4 C)). Neovascularization-related genes mRNA levels were comparable between monoculture (MC) and foetal neural stem (FNS) cells in co-culture with microvessels (hCMEC/D3, HUVECs or HMVECs-L cells), that we defined as control conditions, white in heatmaps (Fig 4 C)) and Fig S7 for unprocessed and non-normalized data).

Most of the neovascularization-related genes mRNA levels increased when endothelial cells are co-cultured with GSCs in cell types studied. Ang1 and Ang2 are modulators of endothelial permeability and barrier function *via* Tie2 ^59–62^. In all cases, Ang1 was downregulated, suggesting disruption of vascular stability. Inversely, Ang2 was upregulated to various levels in all samples. The slight upregulation of Tie2 expression in endothelial cells in the microfluidic system might be associated with Ang2-dependent Tie2 induction. VEGFR1 and VEGFR2 was detectable in all samples. This is consistent with previous report for endothelial cells of the tumour neovasculature and in normal brain vessel adjacent to tumours ^63^. Moreover, MMP-2 and MMP-9 were both upregulated in HUVEC. Production of matrix-degrading proteases, particularly MMPs, by endothelial cells is a critical event during angiogenesis ^64^.

Cross-comparing the heat map patterns for the three endothelial cells, we also found that the response of endothelial cells to the co-culture was GSC-specific. The analysis suggested that HUVECs might be more sensitive to factors released by GSCs than hCMEC/D3 cells and HMVECs-L. These findings reflect the nature of the endothelial cell types described by Uwamori et al., where the authors reported that HUVECs had greater potential than brain microvessel endothelial cells in terms of vascular formation^53^. Moreover, a study described by Guarnaccia et al. , showed that the angiogenic properties of endothelial cells in brain tumour of different grades vary, therefore further confirming the necessity for biologically relevant cell models.^65^

#### Cancer cell dynamics studied by live-cell imaging

An important feature of 3D model systems is that they can be designed to fit to be used for microscopy. Here we coupled a 3D co-culture system with live-cell time lapse imaging to track cells behaviour in specific microvessel types (Fig 5 A)). GFP-tagged patient-derived GBM cells (G166) were used to observe their behaviour in an artificial vascular niche. In every 3D device, we monitored the behaviour of 5-15 cancer cells, in a field of view localized in the interface between the outermost side channel and collagen I-based hydrogel chamber. The cancer cell speed (micron/min) in the 3D device with the endothelial cells move faster when compared to cancer cells in the microfluidic device without endothelial cells forming a microvessel (Fig 5 B)). Between the three organ-specific endothelial cells, the presence of brain hCMEC/D3 as microvessel induced the fastest movement of cancer cells. This might indicate that hCMEC/D3 secrete biochemical molecules or express receptors that may activate glioma stem-like cells more than HUVECs or HMVECs-L cells.

**Fig 5:**
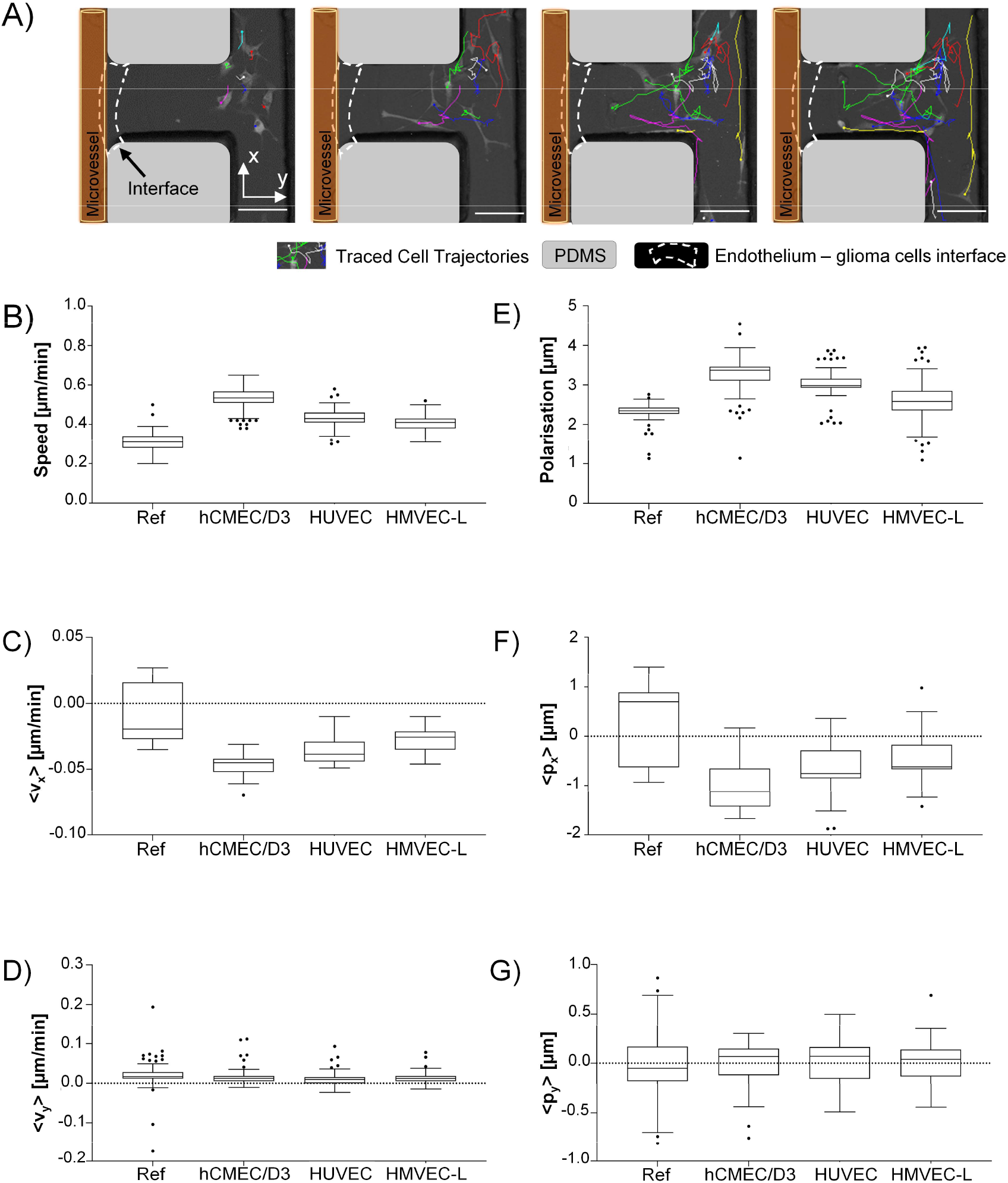
GSCs cultured in the presence of a hCMEC/D3 microvessel are characterised by higher speed, velocity, and preferential polarisation in the direction towards the microvessel. A) Sample images showing tracking of the cell movement in 3-hour time intervals. B) Speed of the cancer cells for the reference and microvascular systems; C) Cancer cell velocity component along the x-axis; D) Cancer cell velocity component along the y-axis; E) Polarisation magnitude of the cancer cells in the collagen matrix; F) Cellular polarisation component along the x-axis; G) Cellular polarisation component along the y-axis. x-axis is situated perpendicular to the barrier, pointing away from the barrier, and y-axis is along the collagen I gel chamber. Scale bar=100 μm. Note that ‘Ref’ samples are measured without any microvessel formation in the side channel.

We also measured to see if the entire cancer cell population (in the FOV) had a preferred direction of migration towards the endothelial cells (Right to left). The velocity components perpendicular to the ECM-endothelium barrier (x axis as shown in Fig 5 A)) and along the ECM chamber were analysed (Fig 5 C)). Cancer cells in the device with a microvessel were shown to move towards the channel interface direction more (i.e. with more negative *x*-velocity component) compared to the microvessel-free control (Fig 5 C)); but no difference was detected in the y-direction (Fig 5 D)). Comparing the data among the three microvessel types, a slight enhancement in directed migration was observed for the cancer cells facing the hCMEC/D3 vessels.

Cancer cell shape axial asymmetries were also investigated during cellular migration inside the collagen I matrix. Brain tumour cells in the presence of hCMEC/D3 microvessel were characterized by the highest median of polarization parameter, as a scalar quantity (Fig 5 E)). As illustrated in Fig 5 F)-G), investigation of cell polarization in x and y directions further indicated that cancer cells appear to favour shifting their mass centre towards the microvessel.

#### Microvessel permeability in co-culture

The permeability of the microvessel serves as an indicator for the endothelial barrier changes due to the presence of tumour in the system. Here, we measured the changes in permeability of vessels formed by the three organ-specific endothelial cell types cultured in 3D ECM microfluidic devices in presence of glioma stem-like cells (G166 or GCGR-E13) or collagen I gel only.

We used the G166 cancer cells, as we assessed the cell movement using this cell line, and the GCGR-E13 cells because of previous characterization for their stemness-related gene expression in 3D ECM culture. Permeability of all three types of endothelium increased in the co-culture setup in comparison to monoculture (collagen I only), as shown in Fig *6* A) i) and B) i). The permeability of endothelium under monoculture conditions is shown in Fig S8.

**Fig 6:**
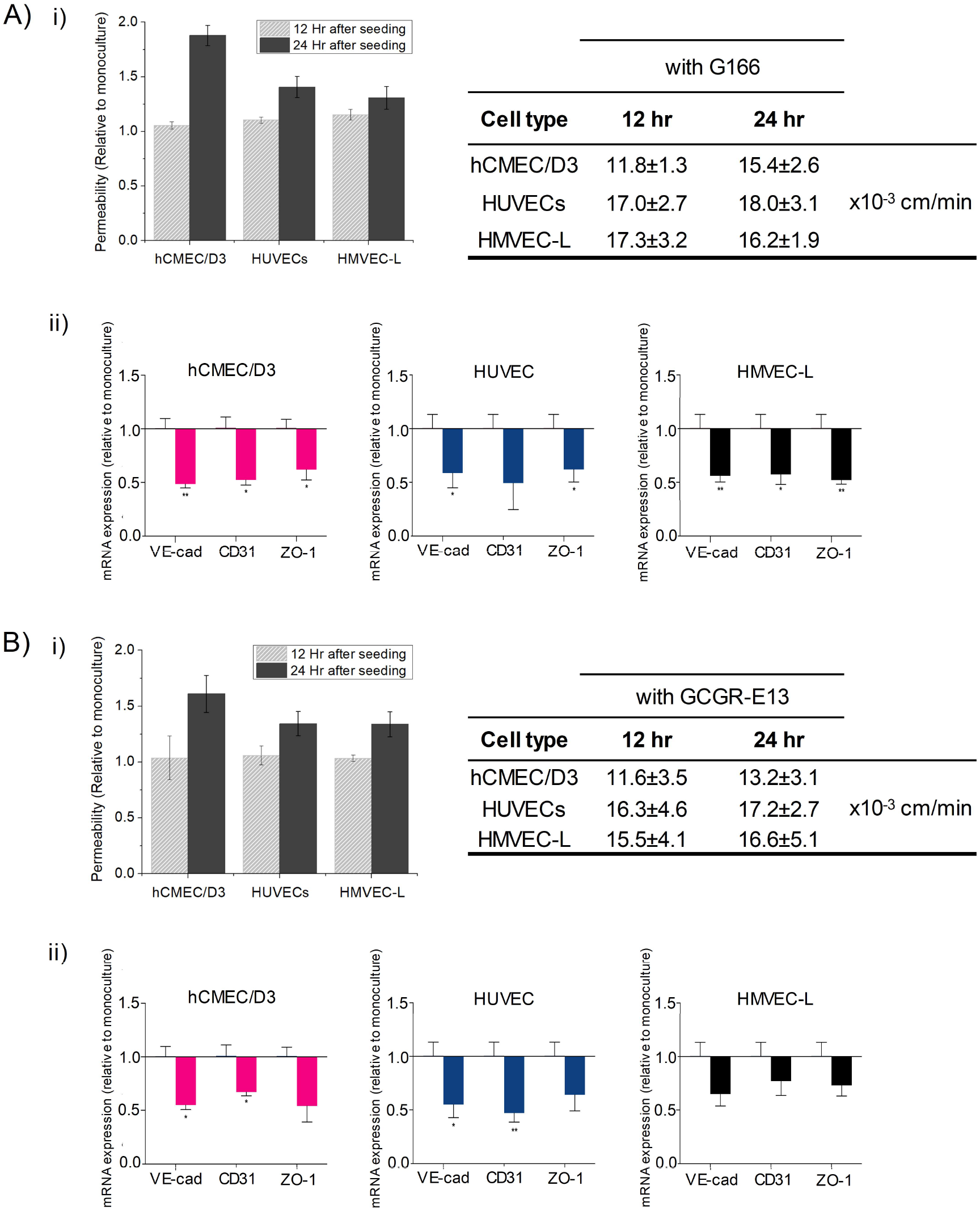
Increased endothelial permeability and the downregulation in mRNA expressions for junction proteins VE-cad, CD31, and ZO-1 in co-cultures with A) G166 and B) GCGR-E13 cancer cells. i) There is an increase in endothelial permeability in co-cultures with G166 and GCGR-E13 cancer cells. The graph shows results from three independent experimental runs as mean ± SEM. Permeability increase correlates with downregulation of tightness-related genes in vessels formed by different vessel types; ii)mRNA expression normalised to samples of endothelial cells cultured without brain cancer cells; results obtained from three independent experimental runs are presented as mean ± SEM. Two-tailed t-test was used for significance.

Several mechanisms exist which could explain changes in permeability due to tumour cell interactions; the permeability increase may be due to tumour cells locally disrupting endothelial monolayer by contact ^66–68^ or through secretion of chemical factors which then compromises the endothelial barrier function ^69^. At the early stages, no cell-endothelium contact was observed during live-cell imaging in the microfluidic system presented here. Thus, we hypothesize that the changes in endothelial permeability might be due to secretion of certain cytokines, such as VEGF or Ang2, by glioblastoma stem-like cells, or endothelial cells themselves ^3,70–72^. On the other hand, no endothelial sprouting was observed even after 3 or 7 days of co-culture, which could suggest different factors contributing to increased vessel permeability.

Moreover, as mentioned earlier, VEGF and Ang2, though mainly associated with pro-angiogenic pathways, have been shown to be present in vessel co-option areas at early stages of GBM formation. It would be necessary to analyse the secretome, as well as the cargo carried in extracellular vesicles that are released by given glioblastoma stem-like cells, to understand what triggers changes in permeability of endothelial cells in co-culture. Additionally, identifying receptors on endothelial cells for released by cancer cells factors would be needed to fully understand the processes.

Since permeability in all types of microvessel was proved to be increased in the presence of cancer cells in the microfluidic devices, it was also of interest to learn how that corresponded to disruption in junction connections between endothelial cells in artificial microvessels. Therefore, after 24 hr in culture, RNA was isolated from endothelial cells recovered from microfluidic chips cultured either without or with GBM cells. For consistency, G166 and GCGR-E13 cells were used to conduct those experiments. Analysis of mRNA levels of *VE-cad*, *CD31* and *ZO-1* revealed downregulation of those genes in all three types of endothelial cells in the co-culture setup (Fig *6* A) ii) and B) ii)). Further to permeability studies, this can indicate disruption in tightness of the microvessels caused by paracrine signalling from cancer cells or autocrine signalling from the endothelial cells themselves, which could be the first step to changes in normal brain vasculature.

## 4 Conclusions

To summarize, the interactions between patient-derived GSC cell lines and microvessels generated from endothelial cells hCMEC/D3, HUVECs and HMVEC-L were studied utilizing a microvessel-on-chip platform. A gravity-driven flow perfusion system was used to maintain endothelium-cancer cell co-culture, and protocols for cell retrieval and live-cell imaging were optimized, thus creating a microfluidic-based assay which could be used for studying GSCs and microvessel interaction. Within a 3D culture, hCMEC/D3 was observed to form a tighter, and therefore more favourable artificial microvessel than HUVECs or HMVECs-L. It was also shown that endothelial cells grown in the serum-free medium created a tubular structure in the microfluidic devices and the formed layer was characterized by tight junction expression and was acceptable for artificial microvessel permeability; thus, overcoming the drawback of using two different media in one experimental setup.

Having optimized the microfluidic platform for GSC-microvessel co-culture, it was found that different cell types (GSCs, U87, foetal neural stem cells) influence gene expression patterns in endothelial cells of three tissue origins, in that various signalling pathways might be involved in initial neovascularization processes depending on brain cell type. Moreover, live-cell imaging and image analysis revealed that brain tumour cells are characterized by increased speed and velocity in the direction of endothelial lining. Additionally, analysis on cell polarization further confirmed asymmetric shift in cell bodies direction towards the vessel. Noticeably, brain tumour cells cultured in the presence of hCMEC/D3 migrated with a higher speed towards the endothelial barrier than in the reference or other microvessel systems. The results therefore proposed that brain-specific microvessel might release certain factors which chemoattract brain tumour cells.

The presented 3D co-culture microsystem, optimized for both endpoint analysis and live-cell imaging, would be beneficial for potential studies of vessel co-option. This proposed microfluidic model might be used for high-throughput culture of various patient-derived GBM cells and to monitor their behaviour in an artificial vascular niche, in highly controllable environment. Further, quantifying parameters, like molecular markers of GBM cells, their speed, polarization, and influence on endothelium, might provide answers on relation between, for instance, specific genetic and epigenetic alterations of GBM cells and the mechanisms of neovascularization they utilize.

The optimisation of the customizable microvessel-on-a-chip device provided additional complexity for in vitro systems and the opportunity to study the influence of glioma cells on normal brain endothelium. Our microfluidic protocol for forming in vitro models of patient-specific glioblastoma-vascular niches, has demonstrated the possibilities to conduct comparative studies to dissect the influence of 3D culture, microvessel architecture and organotypic vessel types on glioma cells' stemness and migration. The possibility to create various customized tumour microenvironments (e.g., choice of ECM components and stiffness, microvessel size and flow rate) might allow clinicians to have the tool to study the tumour responses in differing microenvironment cues, which could lead to swift testing of new drugs and therapeutic approaches to expand GBM treatment options.

## 6 Acknowledgments

This work was supported by the Engineering and Physical Sciences Research Council (EPSRC, EP/S009000/1), European Research Council (ERC-StG 758865, ERC advanced grant REP-789054-1), National Institute for Health Research (NIHR), Cancer Research UK (CRUK), Academy of Medical Sciences, and Cambridge NIHR BRC. M.G. was a recipient of the CRUK Multidisciplinary Studentship. H.B. was a recipient of the CRUK Innovator Award. The GBM cell lines were obtained from the Glioma Cellular Genetics Resource (www.gcgr.org.uk) funded by Cancer Research UK. We thank Prof. Steve Pollard in facilitating the materials transfer. We would also like to thank Charles Cullen for his assistance in establishing the gravity-driven flow for the experiments and Prof. Anne Ridley for advice on the endothelial cell culture.

## 8 Supplementary Materials

**Fig S1:**
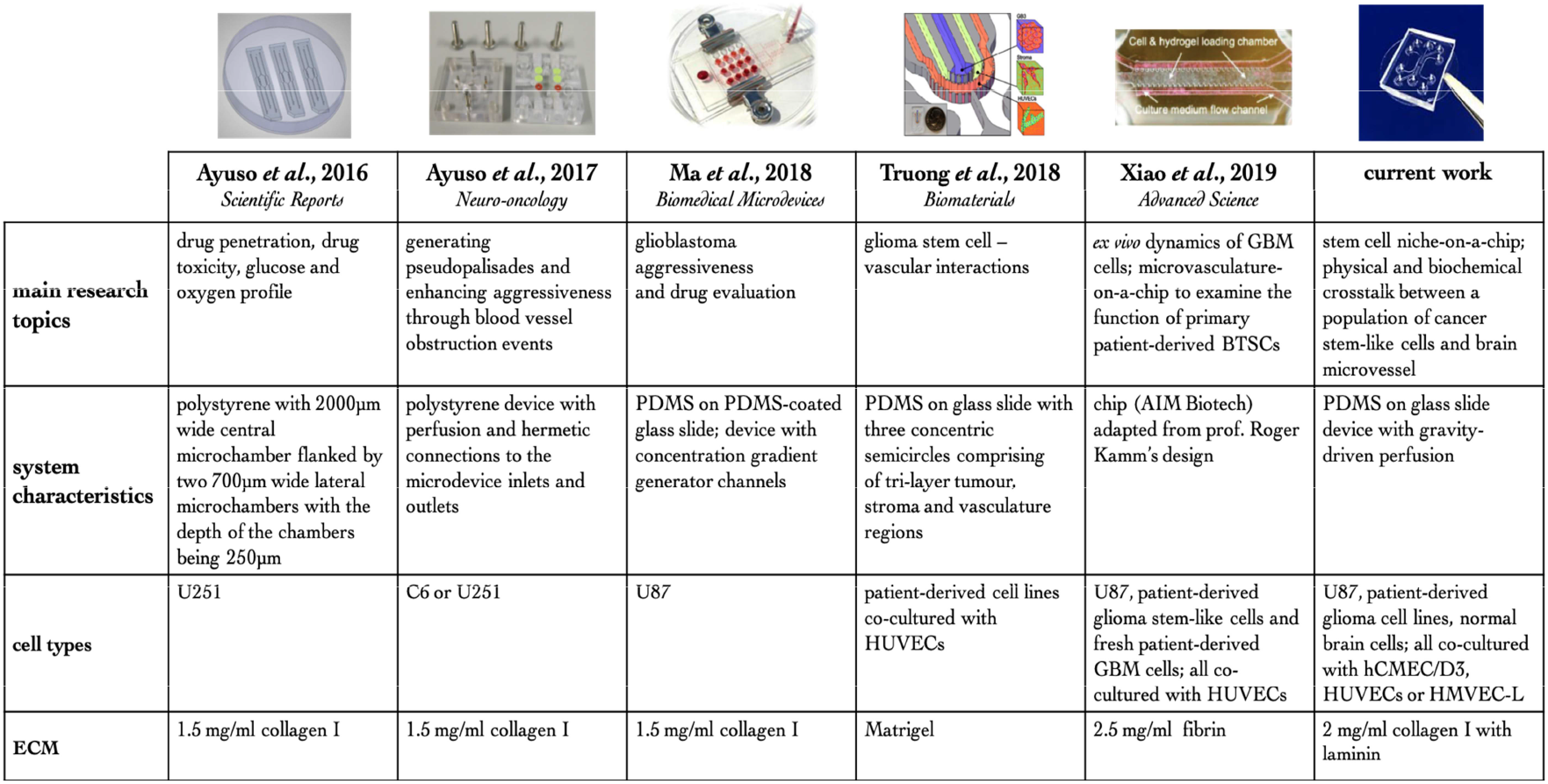
Table summarizing selected microfluidic-based models developed to study GBM cells, or their interactions with vasculature. This table compares published GBM-on-a-chip models to the present work.

**Fig S2:**
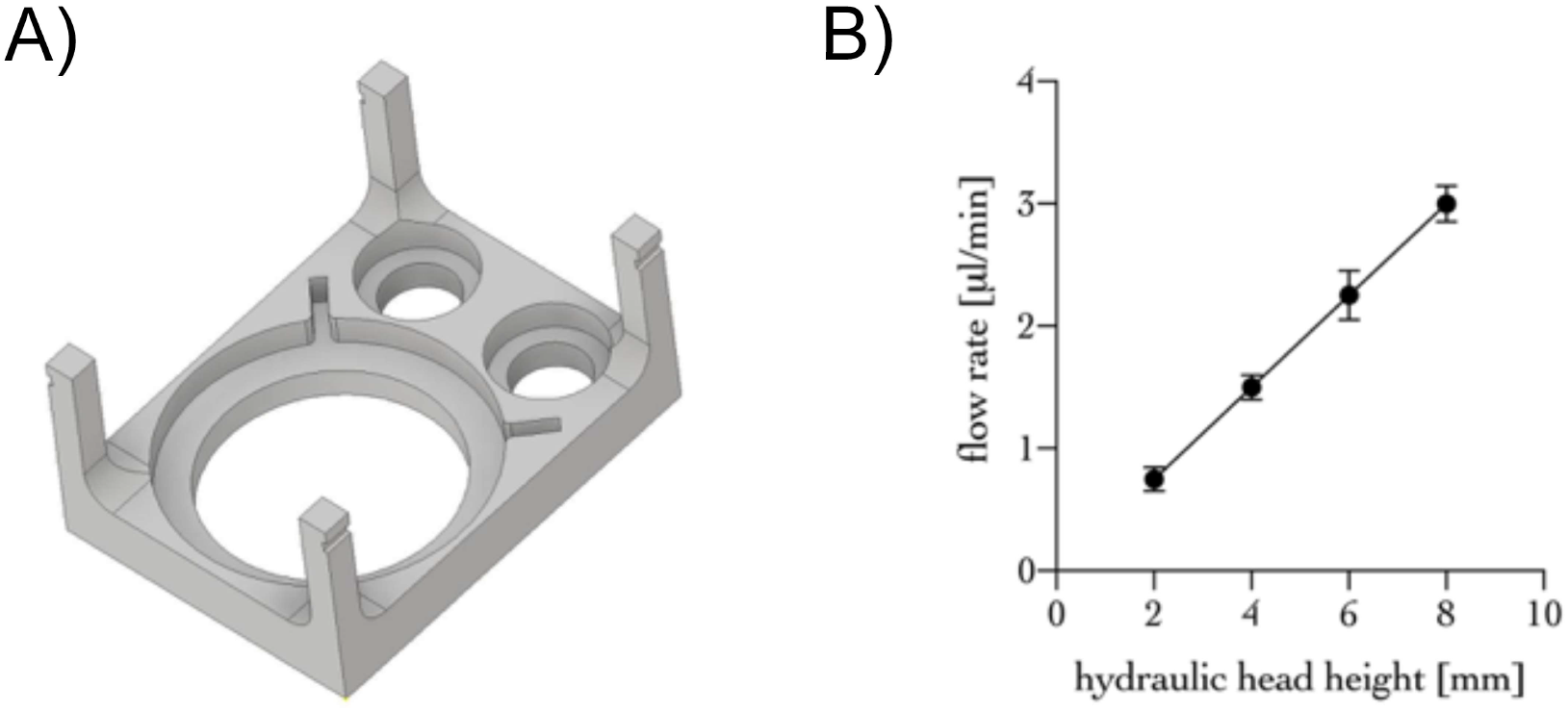
Gravity-driven flow setup. A) design of the gravity-driven pump stage which allows two microfluidic devices and the pump reservoir to be connected easily and safely; B) flow rates obtained from four different hydraulic head heights in the microfluidic device. The graph represents data obtained from three independent experimental runs presented by mean±SEM;

**Fig S3:**
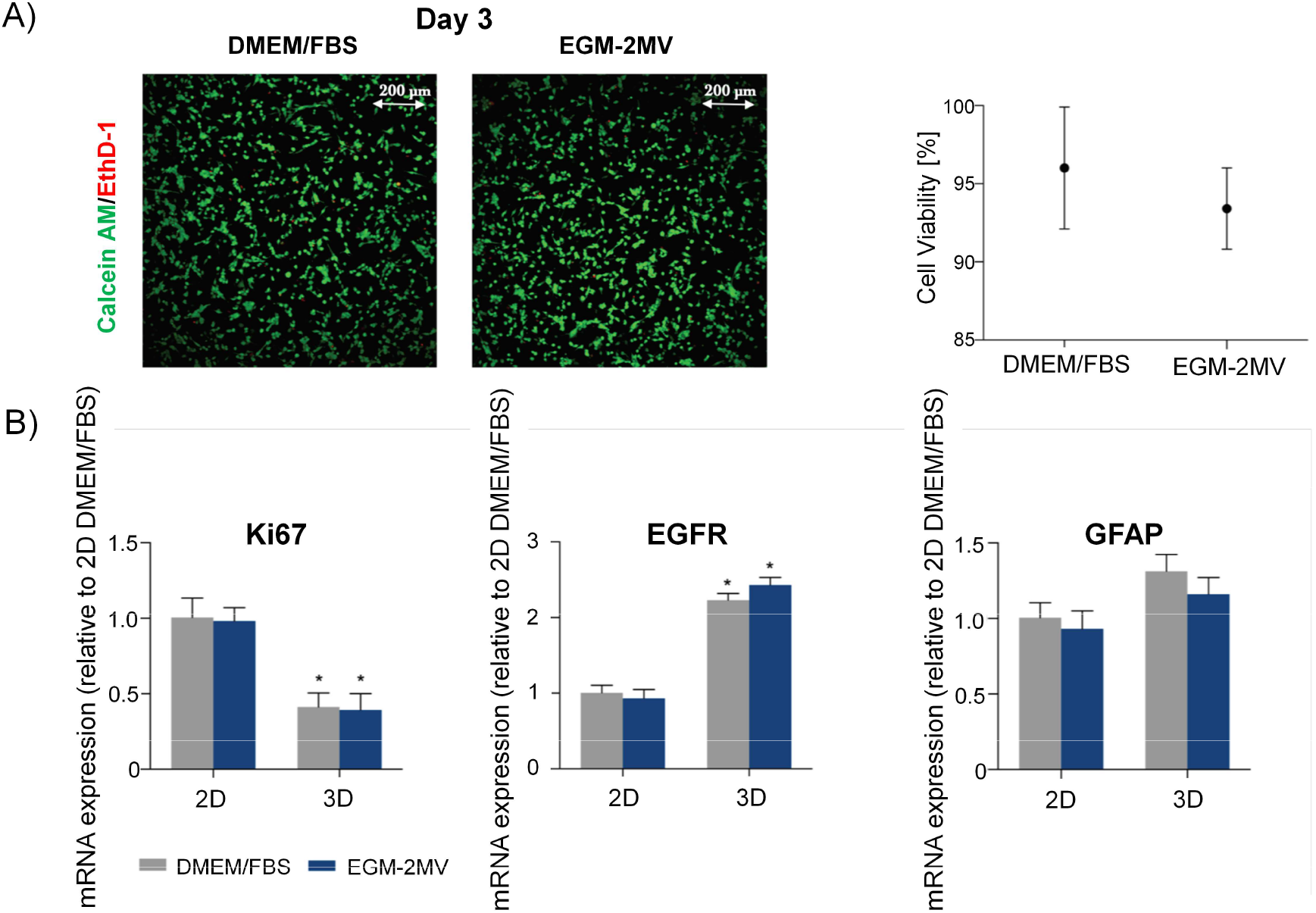
U87 cells can survive in both DMEM/10% FBS and EGM-2MV media whilst cultured in 3D collagen I-based hydrogels in u-slides. A) left side: example images of calcein AM/EthD-1 staining of U87 cells at 3 days in culture; right side: graph presenting viability of U87 cells. Data presented as mean±SD (percentage of live cells); B) mRNA expression normalised to 2D DMEM/10%FBS samples. Results obtained from three independent experimental runs are presented by mean±SEM. One-way ANOVA with Tukey’s post-hoc test was used for significance

**Fig S4:**
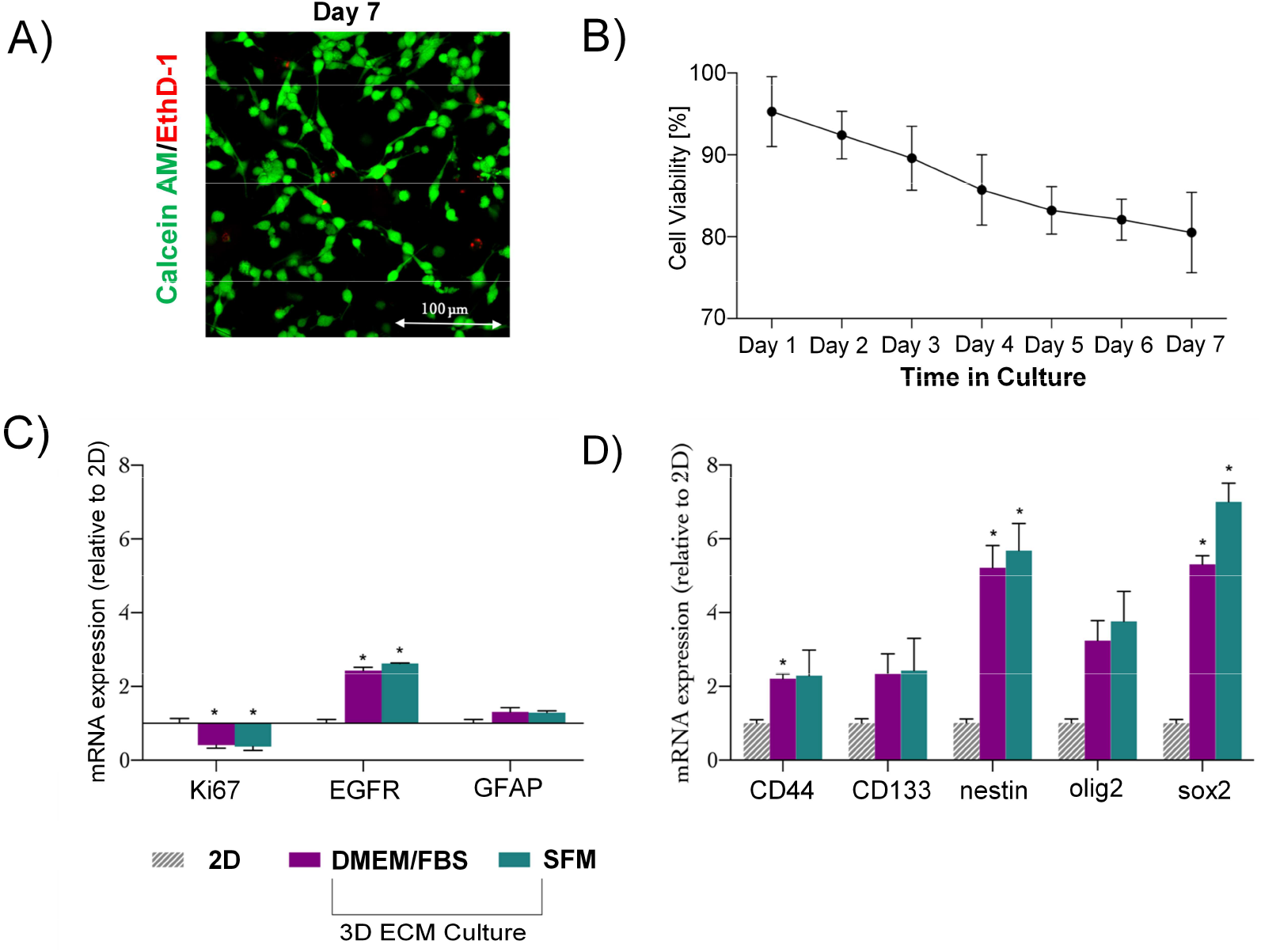
U87 cells cultured in 3D collagen I-based ECM with serum-free medium express increased stemness-related genes compared to 2D culture. (A) Representative image of calcein AM/EthD-1 staining of U87 cells at day-7 culture; (B) Viability of U87 cells over seven days in 3D cell culture. Data presented as mean±SD (percentage of live cells). (C) Ki67 EGRF and GFAP, and (D) mRNA expression of U87 cells grown in 3D collagen I-based gels versus 2D culture for three days (GAPDH was used as housekeeping gene). Results are from three independent experimental runs are presented by mean±SEM. One-way ANOVA with Tukey’s post-hoc test was used for significance.

**Fig S5:**
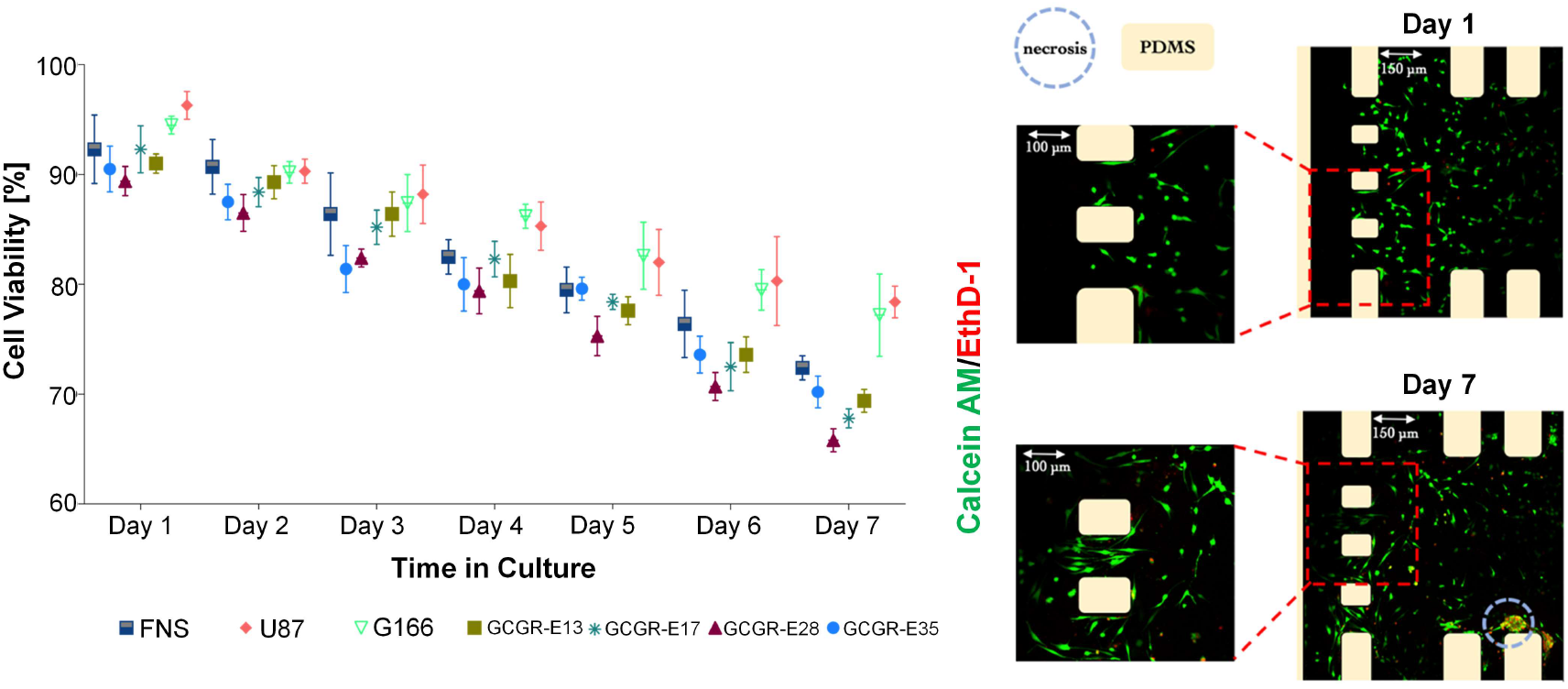
Left: graph presenting viability of six different GSCs and one foetal neural stem cell line over seven days in cell culture; Right: Example images of calcein AM/EthD-1 staining of U87 cells at day 1 and day 7 in culture. At day seven in culture, cells migrated into the outermost side channels, and formed necrotic clusters in the centre of the chamber. The graph shows percentage viability values – data is presented as mean±SD, obtained from three independent experimental runs.

**Fig S6:**
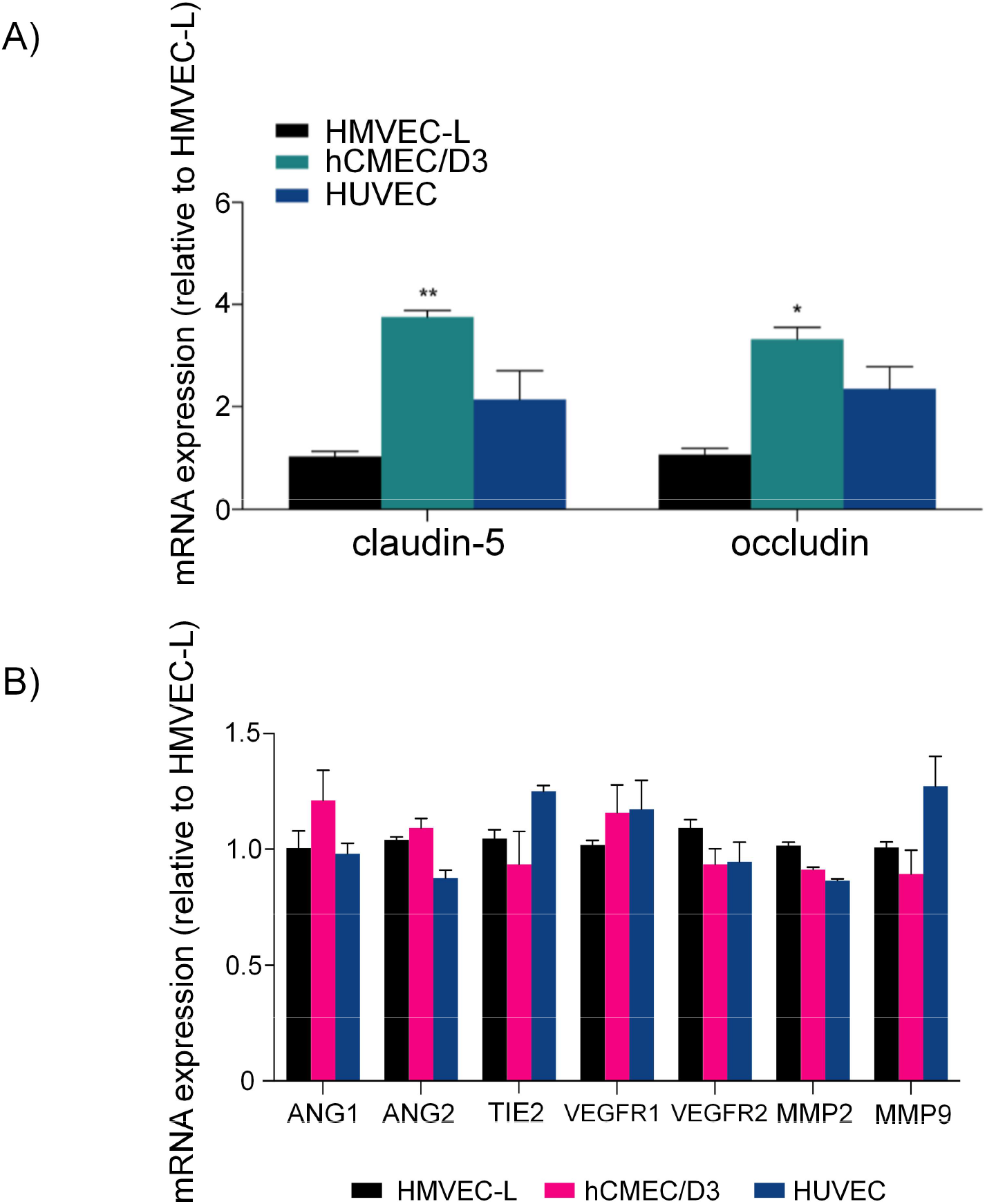
(A) mRNA expression normalised to HMVEC-L as sample of the lowest expression of all tested genes. Results obtained from three independent experimental runs are presented by mean±SEM. One-way ANOVA with Tukey’s post-hoc test was used for significance; (B) In monoculture of microvessels, endothelial cells of different tissue origin are characterised by similar levels of expression of neovascularisation-related genes. mRNA expression normalised to HMVEC-L microvessel. Results obtained from three independent experimental runs are presented by mean±SEM.

**Fig S7:**
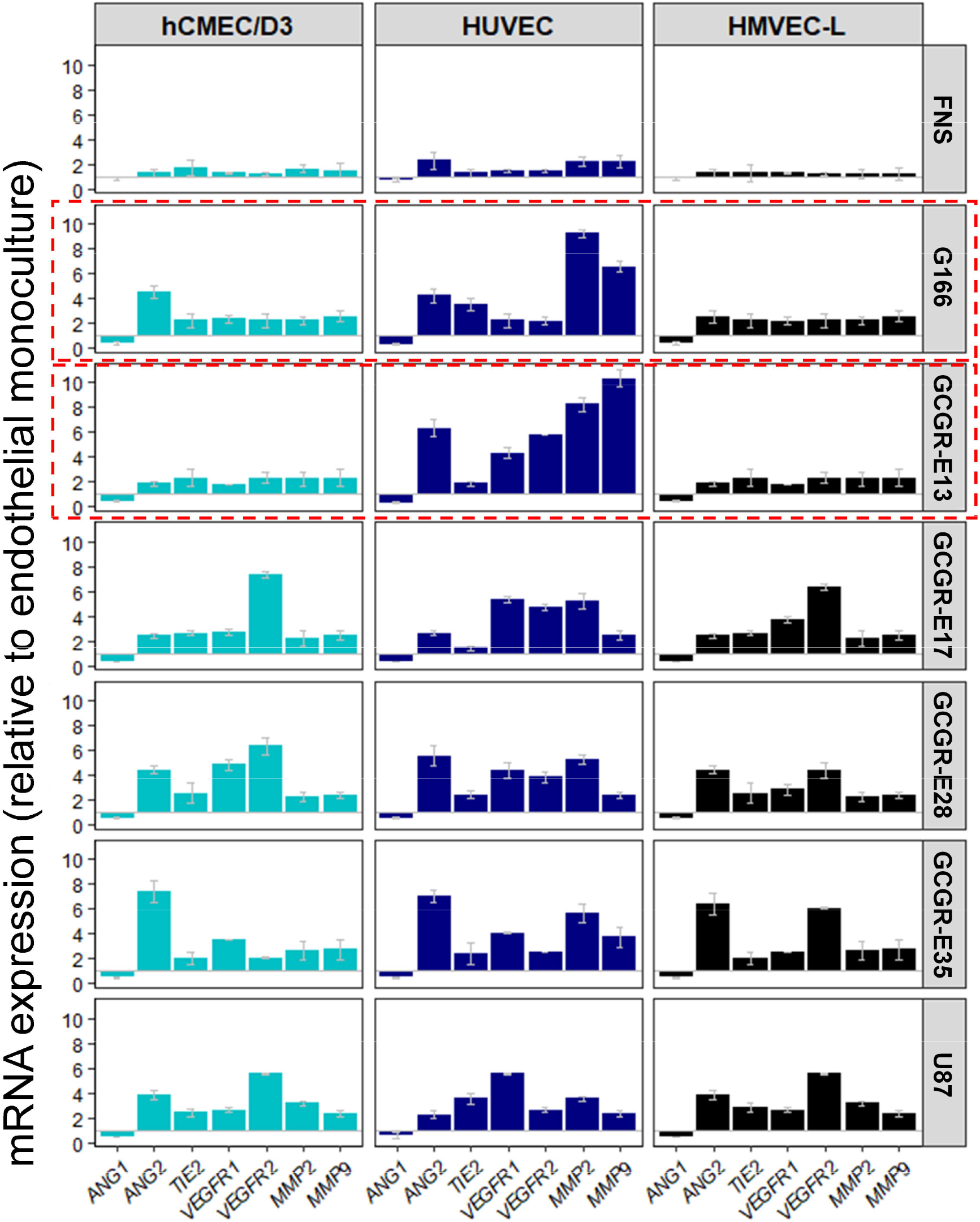
Gene expression in endothelial cells changes under the influence of brain cancer cells in organ-specific manner. Graphs were plotted in RStudio after the analysis of mRNA expression normalised to endothelial monoculture. Results obtained from three independent experimental runs are presented by mean±SEM.

**Fig S8:**
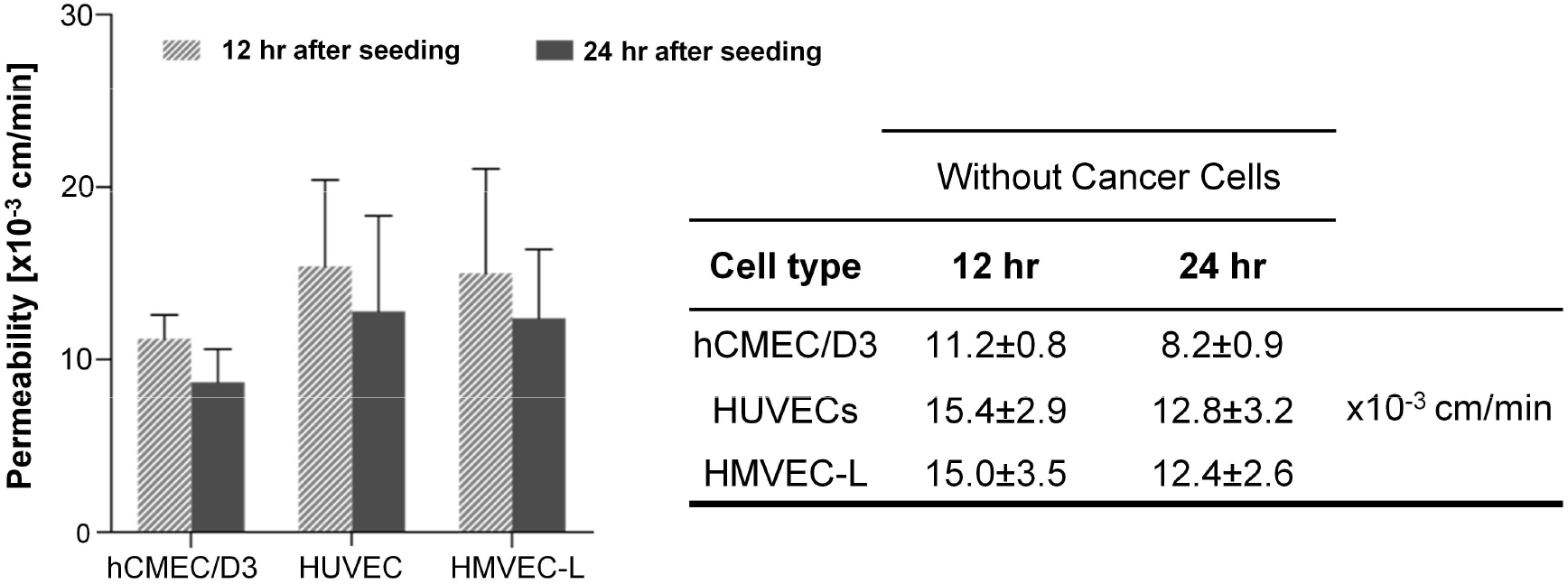
Changes in microvessel permeability without cancer cells after 12 hr and 24 hr. The graph shows results from three independent experimental runs as mean ±SEM.

**Table S1 A):** Primary antibodies.

| **Antigen** | **Host** | **Dilution IF** | **Dilution WB** | **Supplier** |
| --- | --- | --- | --- | --- |
| *CD31* | mouse | 1:200 | 1:1000 | Invitrogen |
| *CD44* | rabbit | 1:500 | 1:2000 | abcam |
| *CD133* | rabbit | 1:200 | 1:500 | abcam |
| *GAPDH* | goat | - | 1:1000 | abcam |
| *GFAP* | mouse | - | 1:1000 | Sigma |
| *Ki67* | rabbit | - | 1:500 | abcam |
| *nestin* | mouse | 1:500 | 1:1000 | Millipore |
| *olig2* | mouse | 1:500 | 1:1000 | Invitrogen |
| *sox2* | mouse | 1:200 | 1:500 | R&D Systems |
| *VE-cad* | mouse | 1:200 | 1:100 | Novus Biologicals |
| *ZO-1* | rabbit | 1:500 | 1:1000 | abcam |

**Table S1 B):** Secondary antibodies and fluorescent probes

| **Secondary Antibodies** |  |  |  |  |
| --- | --- | --- | --- | --- |
| **Antigen** | **Host** | **Conjugation** | **Dilution** | **Supplier** |
| *goat IgG* | donkey | HRP | 1:5000 | R&D Systems |
| *mouse IgG* | sheep | HRP | 1:5000 | R&D Systems |
| *rabbit IgG* | donkey | HRP | 1:5000 | GE Healthcare |
| *mouse IgG* | donkey | Alexa Fluor® 488 | 1:1000 | abcam |
| *rabbit IgG* | goat | Alexa Fluor® 488 | 1:1000 | abcam |
| *mouse IgG* | goat | Alexa Fluor® 647 | 1:1000 | abcam |
| *rabbit IgG* | donkey | Alexa Fluor® 647 | 1:1000 | abcam |
| **Fluorescent probes** |  |  |  |  |
| **Names** |  | **Dilution** | **Supplier** |  |
| *Alexa Fluor 488® phalloidin* |  | 1:5000 | Invitrogen |  |
| *CyTRAK Orange™* |  | 1:1000 | abcam |  |

## References

1. R. M. Quaresma, M. P. Coleman and B. Rachet, Lancet, 2015, 385, 1206–1218.

2. A. Roos, Z. Ding, J. C. Loftus and N. L. Tran, Front. Oncol., 2017, 7, 1.

3. R. K. Jain, Cancer Cell, 2014, 26, 605–622.

4. M. E. Hardee and D. Zagzag, Am. J. Pathol., 2012, 181, 1126–1141.

5. T. Donnem, A. R. Reynolds, E. A. Kuczynski, K. Gatter, P. B. Vermeulen, R. S. Kerbel, A. L. Harris and F. Pezzella, Nat. Rev. Cancer, 2018, 18, 323–336.

6. G. Seano and R. K. Jain, Angiogenesis, 2020, 23, 9–16.

7. S. Nath and G. R. Devi, Pharmacol. Ther., 2016, 163, 94–108.

8. F. Weeber, S. N. Ooft, K. K. Dijkstra and E. E. Voest, Cell Chem. Biol., 2017, 24, 1092–1100.

9. J. M. Ayuso, M. Virumbrales-Muñoz, A. Lacueva, P. M. Lanuza, E. Checa-Chavarria, P. Botella, E. Fernández, M. Doblare, S. J. Allison, R. M. Phillips, J. Pardo, L. J. Fernandez and I. Ochoa, Sci. Rep., , DOI:10.1038/srep36086.

10. J. M. Ayuso, R. Monge, A. Martínez-González, M. Virumbrales-Muñoz, G. A. Llamazares, J. Berganzo, A. Hernández-Laín, J. Santolaria, M. Doblaré, C. Hubert, J. N. Rich, P. Sánchez-Gómez, V. M. Pérez-García, I. Ochoa and L. J. Fernández, Neuro. Oncol., 2017, 19, 503–513.

11. J. Ma, N. Li, Y. Wang, L. Wang, W. Wei, L. Shen, Y. Sun, Y. Jiao, W. Chen and J. Liu, Biomed. Microdevices, , DOI:10.1007/s10544-018-0322-4.

12. D. Truong, R. Fiorelli, E. S. Barrientos, E. L. Melendez, N. Sanai, S. Mehta and M. Nikkhah, Biomaterials, 2019, 198, 63–77.

13. Y. Xiao, D. Kim, B. Dura, K. Zhang, R. Yan, H. Li, E. Han, J. Ip, P. Zou, J. Liu, A. T. Chen, A. O. Vortmeyer, J. Zhou and R. Fan, Adv. Sci., , DOI:10.1002/advs.201801531.

14. M. Campisi, Y. Shin, T. Osaki, C. Hajal, V. Chiono and R. D. Kamm, Biomaterials, 2018, 180, 117–129.

15. G. Adriani, D. Ma, A. Pavesi, R. D. Kamm and E. L. K. Goh, Lab Chip, 2017, 17, 448–459.

16. S. I. Ahn, Y. J. Sei, H.-J. Park, J. Kim, Y. Ryu, J. J. Choi, H.-J. Sung, T. J. MacDonald, A. I. Levey and Y. Kim, Nat. Commun., 2020, 11, 1–12.

17. M. S. Choe, J. S. Kim, H. C. Yeo, C. M. Bae, H. J. Han, K. Baek, W. Chang, K. S. Lim, S. P. Yun and I. Shin, FASEB J.

18. S. Bian, M. Repic, Z. Guo, A. Kavirayani, T. Burkard, J. A. Bagley, C. Krauditsch and J. A. Knoblich, Nat. Methods, 2018, 15, 631–639.

19. C. Bertulli, M. Gerigk, N. Piano, Y. Liu, D. Zhang, T. Müller, T. J. Knowles and Y. Y. S. Huang, Sci. Rep., , DOI:10.1038/s41598-018-30776-0.

20. I. J. Huijbers, M. Iravani, S. Popov, D. Robertson, S. Al-Sarraj, C. Jones and C. M. Isacke, PLoS One, 2010, 5, e9808.

21. S. K. Chintala, Z. L. Gokaslan, Y. Go, R. Sawaya, G. L. Nicolson and J. S. Rao, Clin. Exp. Metastasis, 1996, 14, 358–366.

22. A. C. Bellail, S. B. Hunter, D. J. Brat, C. Tan and E. G. Van Meir, Int. J. Biochem. Cell Biol., 2004, 36, 1046–1069.

23. S. M. Pollard, K. Yoshikawa, I. D. Clarke, D. Danovi, S. Stricker, R. Russell, J. Bayani, R. Head, M. Lee and M. Bernstein, Cell Stem Cell, 2009, 4, 568–580.

24. K. Funamoto, D. Yoshino, K. Matsubara, I. K. Zervantonakis, K. Funamoto, M. Nakayama, J. Masamune, Y. Kimura and R. D. Kamm, Integr. Biol., 2017, 9, 529–538.

25. Y. Liu, E. Gill and Y. Y. Shery Huang, Futur. Sci. OA, 2017, 3, FSO173.

26. S. Brem, R. Cotran and J. Folkman, J. Natl. Cancer Inst., 1972, 48, 347–356.

27. D. N. Louis, H. Ohgaki, O. D. Wiestler, W. K. Cavenee, P. C. Burger, A. Jouvet, B. W. Scheithauer and P. Kleihues, Acta Neuropathol., 2007, 114, 97–109.

28. R. Galli, E. Binda, U. Orfanelli, B. Cipelletti, A. Gritti, S. De Vitis, R. Fiocco, C. Foroni, F. Dimeco and A. Vescovi, Cancer Res., 2004, 64, 7011–7021.

29. X. Hong, K. Chedid and S. N. Kalkanis, Int. J. Oncol., 2012, 41, 1693–1700.

30. A. Rape, B. Ananthanarayanan and S. Kumar, Adv. Drug Deliv. Rev., 2014, 79, 172–183.

31. W. Xiao, A. Sohrabi and S. K. Seidlits, Futur. Sci. OA, 2017, 3.

32. D. Lv, S. C. Yu, Y. F. Ping, H. Wu, X. Zhao, H. Zhang, Y. Cui, B. Chen, X. Zhang, J. Dai, X. W. Bian and X. H. Yao, Oncotarget, 2016, 7, 56904–56914.

33. W. Jia, X. Jiang, W. Liu, L. Wang, B. Zhu, H. Zhu, X. Liu, M. Zhong, D. Xie, W. Huang, W. Jia, S. Li, X. Liu, X. Zuo, D. Cheng, J. Dai and C. Ren, Int. J. Oncol., 2018, 52, 1787–1800.

34. M. Virumbrales-Muñoz, J. M. Ayuso, A. Lacueva, T. Randelovic, M. K. Livingston, D. J. Beebe, S. Oliván, D. Pereboom, M. Doblare, L. Fernández and I. Ochoa, Sci. Rep., , DOI:10.1038/s41598-019-42529-8.

35. J. Lee, S. Kotliarova, Y. Kotliarov, A. Li, Q. Su, N. M. Donin, S. Pastorino, B. W. Purow, N. Christopher, W. Zhang, J. K. Park and H. A. Fine, Cancer Cell, 2006, 9, 391–403.

36. B. Borrell, Nature, 2010, 463, 858.

37. P. F. Ledur, G. R. Onzi, H. Zong and G. Lenz, Oncotarget, 2017, 8, 69185.

38. A. Vartanian, S. K. Singh, S. Agnihotri, S. Jalali, K. Burrell, K. D. Aldape and G. Zadeh, Neuro. Oncol., 2014, 16, 1167–75.

39. D. V. Brown, S. S. Stylli, A. H. Kaye and T. Mantamadiotis, in Advances in Experimental Medicine and Biology, Springer New York LLC, 2019, vol. 1139, pp. 1–21.

40. V. Bramanti, D. Tomassoni, M. Avitabile, F. Amenta and R. Avola, Front. Biosci. - Sch., 2010, 2 S, 558–570.

41. S. Bien-Möller, E. Balz, S. Herzog, L. Plantera, S. Vogelgesang, K. Weitmann, C. Seifert, M. A. Fink, S. Marx, A. Bialke, C. Venugopal, S. K. Singh, W. Hoffmann, B. H. Rauch and H. W. S. Schroeder, Stem Cells Int., 2018, 2018, 9628289.

42. G. Cattoretti, M. H. G. Becker, G. Key, M. Duchrow, C. Schlüuter, J. Galle and J. Gerdes, J. Pathol., 1992, 168, 357–363.

43. J. D. Lathia, S. C. Mack, E. E. Mulkearns-Hubert, C. L. L. Valentim and J. N. Rich, Genes Dev., 2015, 29, 1203–1217.

44. D. Garnier, O. Renoult, M. C. Alves-Guerra, F. Paris and C. Pecqueur, Front. Oncol., 2019, 9.

45. G. Bazzoni, O. M. Martínez-Estrada, F. Orsenigo, M. Cordenonsi, S. Citi and E. Dejana, J. Biol. Chem., 2000, 275, 20520–20526.

46. K. Ebnet, C. U. Schulz, M. K. Meyer Zu Brickwedde, G. G. Pendl and D. Vestweber, J. Biol. Chem., 2000, 275, 27979–27988.

47. A. S. Fanning and J. M. Anderson, Ann. N. Y. Acad. Sci., 2009, 1165, 113–120.

48. C. Greene, N. Hanley and M. Campbell, Fluids Barriers CNS, 2019, 16.

49. T. D. Brown, M. Nowak, A. V. Bayles, B. Prabhakarpandian, P. Karande, J. Lahann, M. E. Helgeson and S. Mitragotri, Bioeng. Transl. Med., , DOI:10.1002/btm2.10126.

50. I. K. Zervantonakis, S. K. Hughes-Alford, J. L. Charest, J. S. Condeelis, F. B. Gertler and R. D. Kamm, Proc. Natl. Acad. Sci., 2012, 109, 13515–13520.

51. A. D. Wong and P. C. Searson, Cancer Res., 2014, 74, 4937–4945.

52. J. S. Jeon, I. K. Zervantonakis, S. Chung, R. D. Kamm and J. L. Charest, PLoS One, 2013, 8, e56910.

53. H. Uwamori, Y. Ono, T. Yamashita, K. Arai and R. Sudo, Microvasc. Res., 2019, 122, 60–70.

54. J. van der Valk, K. Bieback, C. Buta, B. Cochrane, W. G. Dirks, J. Fu, J. J. Hickman, C. Hohensee, R. Kolar, M. Liebsch, F. Pistollato, M. Schulz, D. Thieme, T. Weber, J. Wiest, S. Winkler and G. Gstraunthaler, ALTEX, 2018, 35, 99–118.

55. C. A. Gilbert and A. H. Ross, J. Cell. Biochem., 2009, 108, 1031–1038.

56. T. Nitz, T. Eisenblätter, K. Psathaki and H.-J. Galla, Brain Res., 2003, 981, 30–40.

57. B. Andrée, H. Ichanti, S. Kalies, A. Heisterkamp, S. Strauß, P. M. Vogt, A. Haverich and A. Hilfiker, Sci. Rep., 2019, 9, 1–11.

58. S. Hinkel, K. Mattern, A. Dietzel, S. Reichl and C. C. Müller-Goymann, Int. J. Pharm., 2019, 566, 434–444.

59. N. Jones, Z. Master, J. Jones, D. Bouchard, Y. Gunji, H. Sasaki, R. Daly, K. Alitalo and D. J. Dumont, J. Biol. Chem., 1999, 274, 30896–30905.

60. I. Kim, H. G. Kim, J. N. So, J. H. Kim, H. J. Kwak and G. Y. Koh, Circ. Res., 2000, 86, 24–29.

61. H. T. Yuan, E. V. Khankin, S. A. Karumanchi and S. M. Parikh, Mol. Cell. Biol., 2009, 29, 2011–2022.

62. I. Grenga, A. R. Kwilas, R. N. Donahue, B. Farsaci and J. W. Hodge, J. Immunother. Cancer, 2015, 3, 1–11.

63. J. H. Harmey and D. Stefanik, in VEGF and Cancer, 2004, pp. 72–82.

64. G. Taraboletti, S. D’Ascenzo, P. Borsotti, R. Giavazzi, A. Pavan and V. Dolo, Am. J. Pathol., 2002, 160, 673–680.

65. L. Guarnaccia, S. E. Navone, E. Trombetta, C. Cordiglieri, A. Cherubini, F. M. Crisà, P. Rampini, M. Miozzo, L. Fontana, M. Caroli, M. Locatelli, L. Riboni, R. Campanella and G. Marfia, Sci. Rep., 2018, 8, 1–13.

66. Y. H. Li and C. Zhu, Clin. Exp. Metastasis, 1999, 17, 423–429.

67. S. Kumar and V. M. Weaver, Cancer Metastasis Rev., 2009, 28, 113–127.

68. C. T. Mierke, J. Biol. Chem., 2011, 286, 40025–40037.

69. S. M. Weis and D. A. Cheresh, Nature, 2005, 437, 497–504.

70. S. Bao, Q. Wu, R. E. McLendon, Y. Hao, Q. Shi, A. B. Hjelmeland, M. W. Dewhirst, D. D. Bigner and J. N. Rich, Nature, 2006, 444, 756–760.

71. C. Folkins, Y. Shaked, S. Man, T. Tang, C. R. Lee, Z. Zhu, R. M. Hoffman and R. S. Kerbel, Cancer Res., 2009, 69, 7243–7251.

72. L. Treps, R. Perret, S. Edmond, D. Ricard and J. Gavard, J. Extracell. Vesicles, , DOI:10.1080/20013078.2017.1359479.

73. A. Griveau, G. Seano, S. J. Shelton, R. Kupp, A. Jahangiri, K. Obernier, S. Krishnan, O. R. Lindberg, T. J. Yuen and A.-C. Tien, Cancer Cell, 2018, 33, 874–889.

